# The Role of Environmental Stress in Promoting Mutators Through Evolutionary Rescue: Quantitative Predictions

**DOI:** 10.1101/2025.02.11.637742

**Authors:** Marwa Z Tuffaha, Lindi M Wahl

## Abstract

The role of mutation rate in evolutionary rescue has been extensively explored, but little work has investigated how evolutionary rescue can promote mutators, lineages with higher mutation rates. Under complete linkage, we investigate the likelihood of evolutionary rescue on a mutator background that either emerges *de novo* or pre-exists in the population prior to a severe environmental change. If such an evolutionary rescue event occurs, the mutator lineage sweeps into the population, and thus the environmental stress has promoted mutators. Our findings indicate that mutation rate evolution can substantially boost rescue probabilities, but stronger mutators are most effective when the wildtype has a low mutation rate, while their advantage diminishes for higher wildtype mutation rates. Interestingly, at intermediate wildtype mutation rates, emerging mutators can be almost equally likely to sweep no matter how slowly or quickly the environment changes. However, at low wildtype mutation rates, mutators are only likely to sweep for very slow environmental changes due to the sequential nature of necessary mutations for such sweeps to occur. Finally, we show that pre-existing mutators can be significantly more likely to rescue the population compared to the wildtype, provided the wildtype’s mutation rate is relatively low. This research opens new avenues for investigating mutator dynamics in response to environmental stress.

## Introduction

As environmental conditions shift at an unprecedented pace, many populations face existential challenges that threaten their survival (Gonzalez et al. 2013; Alexander et al. 2014). Navigating these challenges through adaptation is crucial to avoid extinction under harsh environmental pressures (Gomulkiewicz and Holt 1995; Carlson et al. 2014), where phenotypic plasticity (Feiner et al. 2021), genetic (Holt and Gomulkiewicz 1997; Orr and Unckless 2008) and epigenetic (O’Dea et al. 2016) variation serve as bases for evolutionary processes that can promote phenotypes capable of thriving in the new environment. These evolutionary challenges are relevant in conservation biology in response to climate change (Jiao et al. 2020; DeFilippo et al. 2022), in the development of drug resistance due to antimicrobial exposure (Woodford and Ellington 2007; Baquero et al. 2021), and in the progression of nascent tumours as they navigate the impacts of harmful passenger mutations (Alejandre et al. 2024).

Evolutionary rescue occurs when a population avoids extinction by adapting fast enough under challenging conditions (Bell 2017; Gonzalez et al. 2013), which can be the outcome we wish to achieve such as in conservation, or what we want to minimize in the case of drug resistance. Rescuing alleles can arise from *de novo* mutations and/or pre-exist in a population through standing variation (Orr and Unckless 2008, 2014). Their success in saving the population depends on the population’s reproductive strategy (Uecker 2017; Garnier et al. 2023), the complex dynamics of community interactions (van Velzen 2023), and whether the rescue phenotype is a single mutation or multiple mutations away from the wildtype (Osmond *et al*. 2020). It has also been shown that potential spatial structure (Tomasini and Peischl 2022), and density-dependent competition can drive local adaptations and influence overall survival (Uecker *et al*. 2014).

Recent modeling studies have shown that the probability of evolutionary rescue increases with mutation rate, especially in rapidly changing environments (Orr and Unckless 2014; Marrec and Bitbol 2020a; Tanaka and Wahl 2022). Nonetheless, while greater genetic diversity is typically advantageous, at very high mutation rates, the burden of deleterious mutations and lethal mutagenesis, in addition to the increased chances of back mutation, may outweigh this benefit and hinder adaptation (Anciaux *et al*. 2019; Sprouffske *et al*. 2018; Hinsch *et al*. 2023). Also, evolutionary rescue models typically consider a fixed wildtype mutation rate (but see Greenspoon and Mideo (2017)), and discuss the impact of whether this fixed rate is low or high (Orr and Unckless 2014; Marrec and Bitbol 2020a; Tanaka and Wahl 2022). However, the evolution of mutation rates is a significant factor that can impact survival positively or negatively (Sniegowski *et al*. 2000; Denamur and Matic 2006; Ferrare and Good 2024).

The rise of mutators, organisms with higher mutation rates than their ancestors, has been observed in experimental populations and natural populations (Sniegowski *et al*. 1997; Shaver *et al*. 2002; Chopra *et al*. 2003; Raynes and Sniegowski 2014; Gifford *et al*. 2023). Many living *E. coli* mutators have been observed with mutation rates as high as 1000-fold higher than typical experimental strains (Chopra *et al*. 2003; Sane *et al*. 2024), while some strains are expected to have 4500-fold higher mutation rates (Sprouffske et al. 2018). By increasing the pool of available mutations in relatively short time frames, mutators may provide a population with a greater chance of finding advantageous adaptations that allow it to thrive in new circumstances (Greenspoon and Mideo 2017). Yet, lower growth rates have been observed at higher mutation rates (Chopra et al. 2003; Gerrish and García-Lerma 2003), and so while the presence of mutators can quicken evolutionary change, it also prompts critical inquiries regarding the long-term stability and fitness of populations that depend on these mechanisms (Raynes and Sniegowski 2014). A particularly important application of evolutionary rescue is the development of anti-microbial resistance, in which mutator strains have been widely observed to develop resistance (i.e., rescue the population and hitchhike) in clinical (Woodford and Ellington 2007; Ramiro et al. 2020; Vanderwoude et al. 2024) and experimental (Chopra et al. 2003; Gifford et al. 2023; Martinez and Baquero 2000) settings. The role of mutators has been found especially crucial in evolving multi-resistance to a combination of drugs (Maciá et al. 2005; Gifford et al. 2023).

Mutation-rate evolution in an evolutionary-rescue setting has so far been explored almost exclusively with simulation frameworks. Romero-Mujalli *et al*. (2019) modelled a diploid sexual population tracking a moving climate optimum and found that mutator alleles invade only under complete linkage or unrealistically high fractions of beneficial mutations. In a medical context, Greenspoon and Mideo (2017) embedded a mutator-modifier locus in a stochastic epidemiological model of parasites exposed to periodic drug switches; mutators could either facilitate or hinder parasite persistence depending on recombination, virulence and the drug-cycling period, but the cost of a high mutation rate was represented by a fixed penalty and no closed-form expressions for rescue likelihood were derived. Most recently, Gifford *et al*. (2023) paired experimental evolution of *E. coli* with a stochastic population-dynamic simulation: their model showed that mutators accelerate the sequential acquisition of multi-drug resistance and can even sweep under single-drug treatment, yet the results again rely on large Monte-Carlo ensembles.

Here we present a mathematically tractable two-locus model of evolutionary rescue with complete linkage between a modifier and a rescue locus, in a continuously deteriorating environment. Our framework (i) yields closed-form approximations for the probability that rescuing lineages arise from either standing or *de novo* mutators, (ii) incorporates a cost function that increases with the absolute mutation rate, thus capturing lethal-mutagenesis limits, and (iii) scales mutation rates at both loci to the genomic mutation rate using experimental estimates. We use this framework to quantitatively explore two closely linked questions: how facing an evolutionary rescue scenario can promote the emergence and fixation of mutator strains, and how mutators can increase the likelihood of surviving such a scenario.

## Methods

Environmental changes can occur in various forms such as abrupt shifts causing sudden fitness loss (e.g., the introduction of a new drug) (Orr and Unckless 2008, 2014; Tanaka and Wahl 2022), gradual changes (e.g., climate change) (Marrec and Bitbol 2020a), periodic fluctuations (e.g., drug cycling) (Greenspoon and Mideo 2017; Marrec and Bitbol 2020b), and unpredictable variations (e.g., resource availability) (Xu et al. 2023; Xu 2023). Gradual environmental changes, relative to the reproductive timescale of the organism, have been shown to improve chances for adaptation and increase the probability of rescue compared to sudden changes (Wu *et al*. 2014), whereas strong stochastic fluctuations in the environment have been proposed to hinder population survival (Xu *et al*. 2023; Marrec and Bank 2023). In scenarios of gradual environmental change, the fitness of wild-type individuals may decline as they become less suited to new conditions, while rescue alleles gradually gain fitness advantages. The shape of the gradual change in fitness can be linear (Chevin *et al*. 2010; Orive *et al*. 2019), but nonlinear forms have been recently suggested (Marrec and Bitbol 2020a; Greenspoon and Spencer 2021) as more suitable models under the current acceleration in climate change (Callaghan *et al*. 2010; Nerem *et al*. 2018).

Following the approach adopted by Marrec and Bitbol (Marrec and Bitbol 2020a), which allows for both gradual and abrupt environmental changes, we consider a wildtype population in a non-linearly deteriorating environment. The authors consider a Hill function to describe the relative growth rate of the wildtype over time 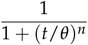, where *n* is the parameter that determines the speed of the environmental change centered around the time *θ*. For convenience, we repeatedly refer to the “relative growth rate” as the “growth rate” throughout this article.

Let *ν >* 0 be the wildtype’s mutation rate per genome per generation. We assume a cost for mutation, *C*(*ν*), that reduces growth rates at higher mutation rates; this is also consistent with empirical fitness measurements in mutator strains (Chopra *et al*. 2003; Gerrish and García-Lerma 2003). We suppose that there is a level at which mutation rates are too high, leading to lethal mutagenesis, by assuming a critical mutation rate, *ν*_*c*_, that causes the growth rate to drop to zero. For simplicity, we assume a linear form for the mutational cost *C*(*ν*) = *ν*/*ν*_*c*_. We thus multiply the growth rate function by max(0, 1 − *C*(*ν*)), where the maximum with zero is taken to avoid negative growth rates. Hence, our growth rate function is given by:

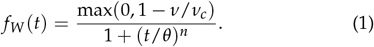

Consider a rescue locus at which the wildtype mutates with rate *µ* per generation and no back mutations are allowed. Once an individual carries the rescue allele, its growth rate is then given by the rising Hill function:

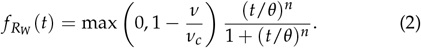

The speed of growth rate decay for the wildtype, determined by *n*, can be different from the speed of the simultaneous increase for the rescue allele. However, throughout the results to follow we consider them equal for simplicity.

We allow another type of mutation at a mutation rate modifier locus, which leads to the emergence of a mutator with an *F*-fold higher mutation rate than the wildtype so that its mutation rate is *ν*^*′*^ = *Fν* per genome per generation. The mutator will have a decaying growth rate function

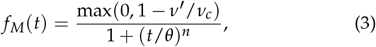

unless a mutation occurs at the rescue locus increasing its growth rate through time as given by the Hill function:

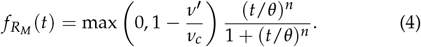

Since the critical value for the mutation rate is a constant, *ν*_*c*_, higher mutation rates for the wildtype, *ν*, correspond to increasing mutation rate costs for both the mutator and wildtype (fig. 1). In other words, multiplying *ν* by the same mutation rate multiple *F* is more deleterious for higher values of *ν*.

**Figure 1.**
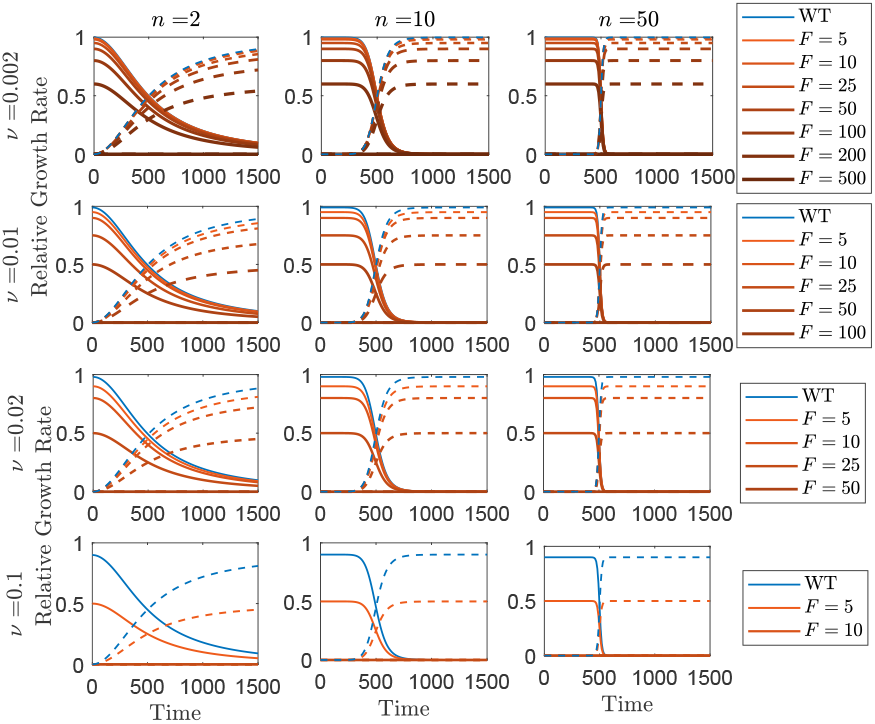
Speed of environmental changes (columns), and wildtype mutation rate, *ν*, (rows), affect the form of the Hill functions for the relative growth rates. Solid lines show growth rate functions for the wildtype (blue) or mutators with different strengths (red) that do not carry the rescue allele, while dashed lines are for the corresponding rescuing mutants. Faster environmental changes (left to right) lead to steeper declines in growth rates around *θ* = 500, while higher initial wildtype mutation rates (top to bottom) restrict the chances for further viable increases in mutation rates (*F* values) as they get closer to the critical mutation rate *ν*_*c*_ = 1.

Following a common terminology in the literature (Travis and Travis 2002; Wylie *et al*. 2009a; Couce *et al*. 2013), we sometimes refer to *F* as the “strength” of the mutator, so that mutators with higher mutation rates are said to be “stronger” in that sense.

### Analytical Predictions

Our analytical approximations of the probabilities of rescue scale all mutation rates to the wildtype mutation rate, allowing us to investigate the lethal mutagenesis limit. As explained in detail in the sections to follow, given the wildtype mutation rate *ν*, the mutator mutation rate is *ν*^*′*^ = *Fν*, while the mutation rate to the rescue allele is *µ* = *rν* for the wildtype and *µ*^*′*^ = *Frν* for the mutator (fig. 2). We consider two ways of introducing mutators into the population (fig. 2). In the first case, mutators can come from *de novo* mutations at a mutation-rate-modifier locus at rate *µ*_*M*_ per generation, such that mutator individuals emerge stochastically from a deterministic wildtype population. The rate *µ*_*M*_ also scales with the wildtype mutation rate (see “Parameter values”). A stochastic approach is required in this case as mutants have a chance of being lost by genetic drift when rare. Alternatively, motivated by recent experimental work (Gifford et al. 2023), mutators can pre-exist at a certain percentage *α* of the population prior to the environmental change. In this case the mutator population declines through time due to both the mutation rate cost and the environmental change, so we consider a deterministic decline for the mutator population. To examine these two cases in isolation, note that there is no *de novo* mutation from wildtype to mutator in the pre-existing mutator case. One can consider the case of having both pre-existence and *de novo* emergence of mutators together, which we leave to future work. In either case, the wildtype and the mutator populations grow logistically and are assumed to have the same death rate *g*, and the total population is always subject to carrying capacity *K*. The rescue allele mutants emerge stochastically from the wildtype and the mutator populations and, for analytical tractability, are presumed to be too rare to impact the carrying capacity of the wildtype or the mutator populations prior to their fixation (we relax this assumption in the simulations). No back mutations are allowed, and the mutators are assumed to have no further increases in mutation rate (they do not mutate at mutation-rate-modifier loci). Although biologically plausible, we do not allow post-rescue mutations at the mutator locus, but we show in the supplement (section S1.3.3 and fig. S7) that this does not affect the results due to the rarity of rescue mutants. All symbols used for variables and parameters are defined in Table 1.

**Table 1.**
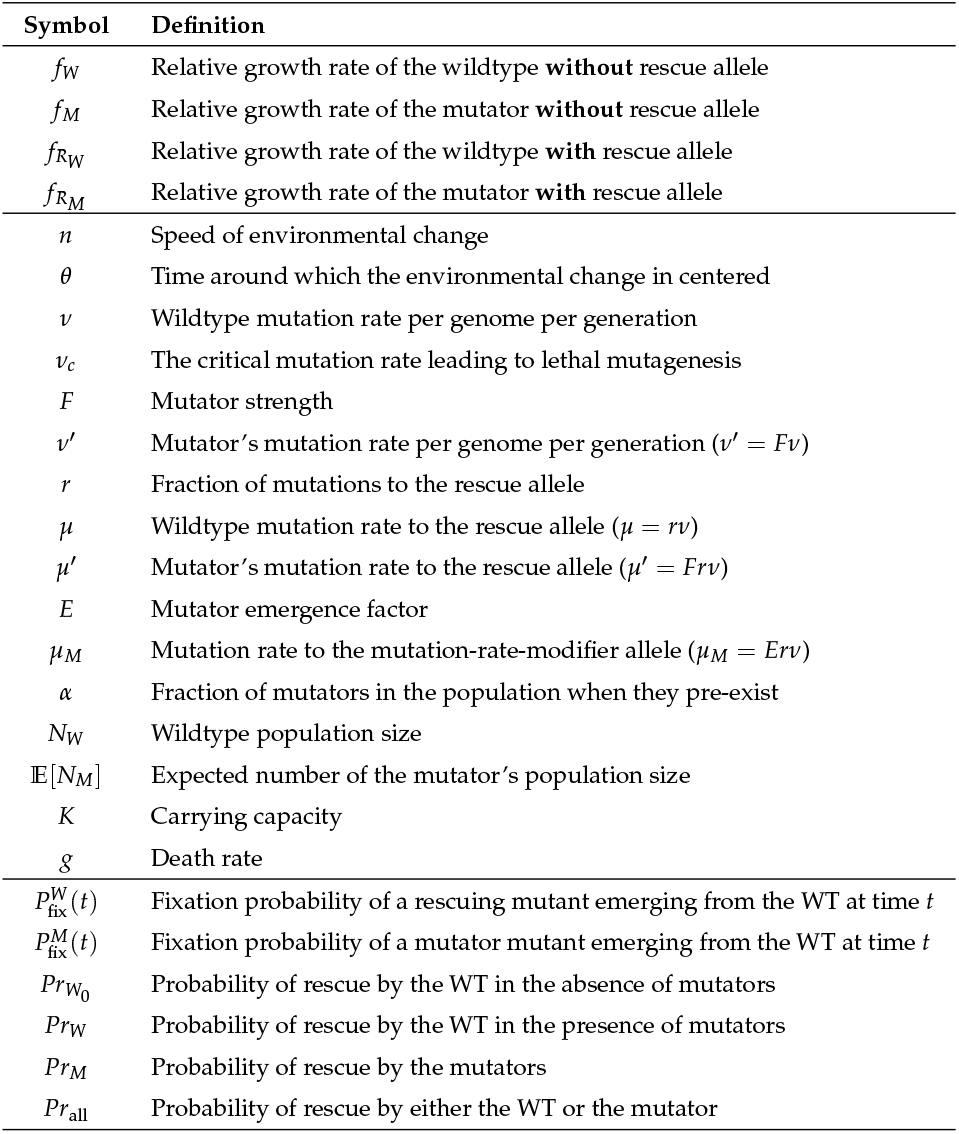
Symbols used in the theoretical analysis. WT=wildtype.

**Figure 2.**
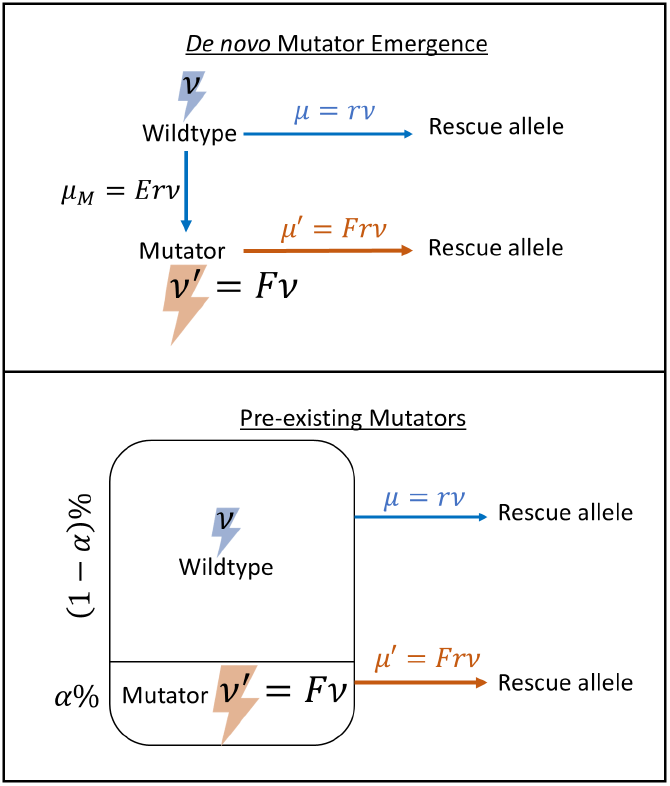
Schematic for modelling evolutionary rescue by mutators. The wildtype’s mutation rate per genome per generation, *ν*, is increased to *ν*^*′*^ = *Fν* for the mutator. Mutators can either emerge from the wildtype population through *de novo* mutations at rate *µ*_*M*_ per generation, or pre-exist at a fraction *α* of the total population. Both the wildtype and mutators mutate at the rescue allele with a fraction *r* of their total possible mutations.

### De novo *Mutator Emergence*

We assume a deterministic decline of the wildtype population governed by the differential equation:

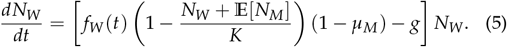

Although we use a stochastic approach to take the effect of genetic drift into account for rare mutators, the mutator population can become large and compete with the wildtype at high emergence rates *µ*_*M*_, and thus we account for competition between the wildtype population and the mutator lineage by including the expected number of mutator individuals, 𝔼[*N*_*M*_], in the carrying capacity term. Since mutator competition is only important for relatively large mutator populations, we use a deterministic approximation for 𝔼[*N*_*M*_] given by:

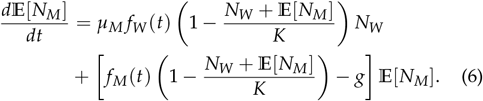

Solving system (5-6) gives an approximation for the wildtype population size at any time.

We note that even for rare mutator populations, the term (1 − *µ*_*M*_) is important in equation (5) as there are cases where the mutator populations stay very small (or even do not exist) even though their emergence rate *µ*_*M*_ is relatively large. One example is when the mutator’s growth rate is very low (or zero) due to the cost of mutations. In this case most emerging mutator individuals do not reproduce, but they nonetheless reduce the wildtype population significantly if *µ*_*M*_ is quite large.

If the emergence rate *µ*_*M*_ is small, then 𝔼[*N*_*M*_] is negligible and the decline of the wildtype population approaches the case of having no mutators, i.e., the dynamics can be well-approximated by removing *µ*_*M*_ and 𝔼[*N*_*M*_] from the equation (5).

We further assume that the mutator’s mutation rate at the rescue locus, *µ*^*′*^ = *Fµ*, is *F*-fold higher than the wildtype’s mutation rate at this locus, *µ*. An individual carrying the rescue allele has a chance of loss by drift. Otherwise, it becomes fixed and rescues the population. Our goal is to find the probability of rescue by the wildtype or by mutators.

### A. Rescue by the Wildtype

In the supplementary material (see S.1.1.1), we follow Marrec and Bitbol (2020a) and use branching processes and a single-variable probability generating function (pgf) to estimate the fixation probability of a rescuing mutant that emerges in a single copy at time *t*_0_,

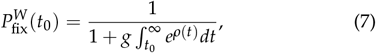

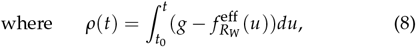

and 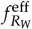 is the effective growth rate of the rescue lineage, as defined in section S.1.

Given 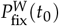, and since the rate at which rescue mutants are generated from the wildtype at time *t* is 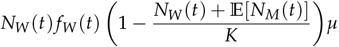, the probability of rescue by the wildtype can be given by:

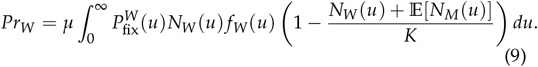

In the results to follow, we will at some points compare the rescue probability by the wildtype in the presence or in the absence of mutators. For the latter case, we set *µ*_*M*_ = 0 (leading to 𝔼[*N*_*M*_ (*u*)] = 0). We then use equation (9) to find the probability of rescue in the absence of mutators, which we denote 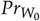.

### B. Rescue by the Mutator

Similarly, we want to estimate the fixation probability of a mutator that emerges as a single copy at time 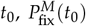. This can be done using a multi-variable probability generating function since fixation of mutators can only occur when a rescue mutant generated by the mutator rescues the population. Here, we do not have an explicit formula as in the case of the wildtype (eq. 7), but instead solve the system numerically (see Supplementary section S.1.1.2).

Once 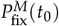 is evaluated, the probability of rescue by the mutator can be given by:

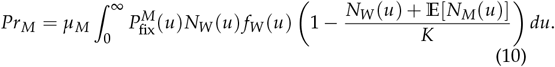

### C. Total Rescue Probability

Once the population has been rescued by either the wildtype or the mutator, it cannot be rescued by the other. This also allows us to calculate the overall probability of rescue by either the wildtype or the mutator as follows:

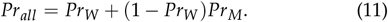

To show the advantage of allowing the evolution of mutators, we define the “mutator advantage” as the relative difference between the overall probability of rescue when mutators are present compared to when they are absent (see Supplementary Section S1.4 for another potential measure):

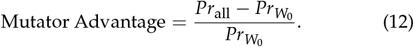

In the above work, we assume a single mutator population with an *F*-fold increase in mutation rate that emerges at rate *µ*_*M*_. In reality, a declining population might have multiple mutators that can emerge and rescue it. This case is modeled in the supplement (S2).

#### Pre-existing Mutator Populations

The impact of mutators on evolutionary dynamics is typically assessed through competition experiments (Sane et al. 2023; Gifford et al. 2023; Thompson et al. 2006). These experiments involve the introduction of mutators as a fraction of the population in order to assess their influence on population outcomes. To model such scenarios, we approximate the probability of rescue by a mutator population that pre-exists prior to the environmental change at fraction *α* of the total population; no *de novo* mutations at the mutation-rate-modifier locus in the wildtype are considered. Two factors cause a decline in the mutator population in this case: (A) the growth rate loss through time due to a worsening environment leads to a decline in both the wildtype and the mutator populations; (B) the mutator always has lower growth rate than the wildtype due to its higher cost of mutation, and thus the mutator population declines even without the environmental change. This second decline can occur rapidly or slowly depending on the difference between the wildtype and mutator growth rates, which can be negligible for low wildtype mutation rates, *ν*, and substantial for high values of *ν* (see fig. 1). These factors ensure an eventual loss of the mutator population, and so we use a deterministic model for the mutator population.

Assume the wildtype and mutator populations, *N*_*W*_ and *N*_*M*_, are governed by the ODE system:

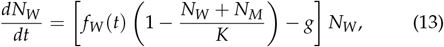

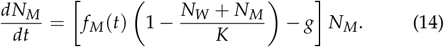

Prior to the environmental change, we assume that the wildtype’s growth rate function is constant and is equal to the its value at time *t* = 0, *f*_*W*_ (0). Thus, if the mutator population does not exist, an existing wildtype population prior to the environmental change stabilizes at 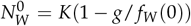. Therefore, we artificially assume this 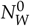 as the total population size even in the presence of the mutator, and impose initial conditions on the above system so that there exists a mutator fraction *α*. Thus, our initial conditions are *N*_*W*_ (0) = (1 − *α*)*K*(1 − *g*/ *f*_*W*_ (0)) and *N*_*M*_ (0) = *αK*(1 − *g*/ *f*_*W*_ (0)). As before, we assume that the wildtype’s mutation rate at the rescue locus, *µ*, increases to *µ*^*′*^ = *Fµ* for the mutator.

We can now use the approximation found earlier by Marrec and Bitbol (2020a) and presented in the previous section for the probability of rescue by mutants founded by deterministic populations. As mentioned earlier, the authors find expressions for a stochastic rescue allele coming from a deterministic wildtype population, which we extend here to have another stochastic rescue population founded by the deterministic mutator.

Following the process described in supplementary section S.1.1.1, but changing the effective growth rate of the rescue mutant to 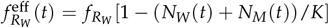, the fixation probability of an individual with the rescue allele emerging from the wildtype at time *t*_0_ is given by:

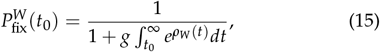

Where

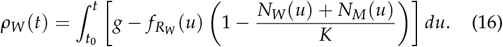

The probability of rescue by the wildtype is given by:

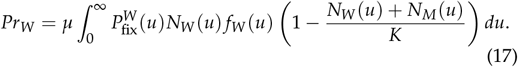

Similarly, the fixation probability of an individual with the rescue allele emerging from the mutator at time *t*_0_ is:

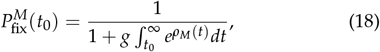

Where

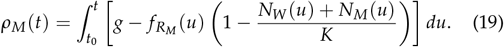

And the probability of rescue by the mutator is:

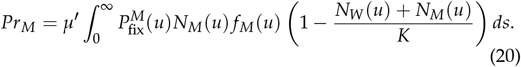

Again, the probability of rescue by either the wildtype or the mutator is *Pr*_*all*_ = *Pr*_*W*_ + (1 − *Pr*_*W*_)*Pr*_*M*_.

#### Expected Advantage of Mutator Pre-existence

By definition, similar to the case of emerging mutators, the advantage of mutator pre-existence can be found using eq. (12), which would now compare the probability of rescue when the population is merely wildtype, 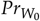, with the overall probability of rescue, *Pr*_all_, when a fraction *α* of the population is initially replaced by mutator individuals.

In fact, if mutation rates had no costs, which is almost the case in this model at very low wildtype mutation rates, we could estimate the advantage of mutator pre-existence without finding any probabilities of rescue. In the absence of mutation cost, introducing mutators only provides more access to mutations at the rescue allele. When *α*% of the population is replaced by mutators that have *F*-fold extra access to the rescue locus mutations, this means we increase the access to these mutations by (*F* − 1)*α*% compared to the case in the absence of mutators. In other words, the effect of this replacement is similar to the effect of increasing the wildtype population size by (*F* − 1)*α*%, which, when survival probabilities are low, should increase the survival chances by the same fraction. We denote this approximate prediction for the mutant advantage as

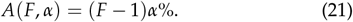

Note: In the results to follow, we show simulation points whenever a probability of rescue is plotted, but not when finding the mutator advantage (eq. (12)). The numerator and denominator become very small at low wildtype mutation rates and this imposes numerical and computational barriers for finding this value by simulation. Thus, we only show analytical predictions for the mutator advantage. Also, for each simulation point, we choose a large enough number of replicates so that the error bars within one standard error are smaller than the symbol size on the results figures, and thus error bars have been omitted throughout. The differences in symbol sizes within figure panels are only for clarity and do not provide any extra information.

#### Parameter Values

In the results to follow, we show the probabilities of rescue for a range of wildtype mutation rates, *ν*, per genome per generation. The values of *ν* start from 10^−4^ per genome per generation, which is the order of magnitude of *E. coli* (Jee et al. 2016), up to a critical mutation rate, *ν*_*c*_. Many living *E. coli* mutators have been observed with mutation rates as high as 0.1 per genome per generation (Sane et al. 2024), so we assume that a mutation rate that is 10-fold higher than that leads to mutational meltdown, that is *ν*_*c*_ = 1; at this mutation rate all wildtype individuals are expected to have a *de novo* mutation at some locus in the genome. While higher mutation rates might be feasible (e.g. Anciaux et al. (2019) use a higher critical value), this conservative choice does not affect our qualitative results. High values of *ν* imply that the wildtype itself is a mutator, and thus further increases in mutation rates are unlikely to offer a benefit, leaving little room for mutator emergence.

We choose a carrying capacity *K* = 10^4^; higher values of *K* would be more realistic for microbial populations but the computational cost of the simulations increases substantially.

For the pre-existing mutators case, we choose a mutator fraction *α* = 5% of the total population, a common percentage used in microbial laboratory experiments (Sane et al. 2024; Gifford et al. 2023).

We further assume that a small fraction, *r*, of all possible mutations occur at the rescue locus, such that the wildtype’s mutation rate at the rescue locus is *µ* = *rν*. Assuming a single gene is responsible for rescue in the E. coli genome, which is approximately 5 Mbp long (Blattner et al. 1997), and considering that typical strains have around 5,000 genes (Lukjancenko *et al*. 2010), the expected fraction of mutations occurring in an average-sized gene would be about 2 *×* 10^−4^. Conservatively assuming that rescue can occur via a loss-of-function mutation in a single gene, and that any mutation in this gene will eliminate function, this implies that *r* = 2 *×* 10^−4^. In other words, out of every 5000 mutations, one mutation is expected to confer evolutionary rescue. If the mutational target for rescue includes many genes, or only a handful of specific loci, obviously higher of lower values of *r* should be considered. In our model, the smaller the value of *r*, the smaller the probability of rescue, which leads to computational challenges; higher values of *r* leave less room for mutator emergence as explained below.

Although there is no concrete measure of the mutation rate at mutation-rate-modifier loci (Vázquez-Mendoza et al. 2024), it is suggested that mutators, with all possible mutation rate elevations, may emerge at a rate up to 10,000 times higher than the rate of observable (rescue) mutations (Vázquez-Mendoza et al. 2024). In our models, we choose the mutator(s) emergence rate to be *E*-fold higher than the mutation rate to the rescue allele, i.e., *µ*_*M*_ = *Eµ*, where we call *E* the “mutator emergence factor” and vary it between 50 and 2,000. Given the value *r* = 2 *×* 10^−4^, it is not realistic to increase *E* beyond 2,000, or else the mutator emergence rate *µ*_*M*_ will approach the total genomic mutation rate *ν*.

The mutator is assumed to have an *F*-fold higher mutation rate than the wildtype, and we vary *F* between 2 and 80. Mutators with higher values of *F* are only reproductive for wildtype with low mutation rates.

Numerical simulations use a Gillespie algorithm (Gillespie 1977) and are explained in detail in the supplemental section S1.1.

## Results

### Environmental Stress Promotes Mutators

While most of the results to follow are obtained analytically, Figure 3 shows some initial simulation results that establish important intuitions about this rescue process (see section S1.3). Critically, in the absence of environmental stress, mutators fail to fix, highlighting that mutator lineages either co-exist at low frequencies (fig. 3A) or go extinct (fig. 3B) if the environment remains hospitable to the wildtype. In both scenarios where mutators emerge *de novo* (panel A) or where they pre-exist (panel B), mutator fixation (red) is never observed in a constant environment, indicating that environmental stress drives the evolutionary success of mutators via rescue events.

**Figure 3.**
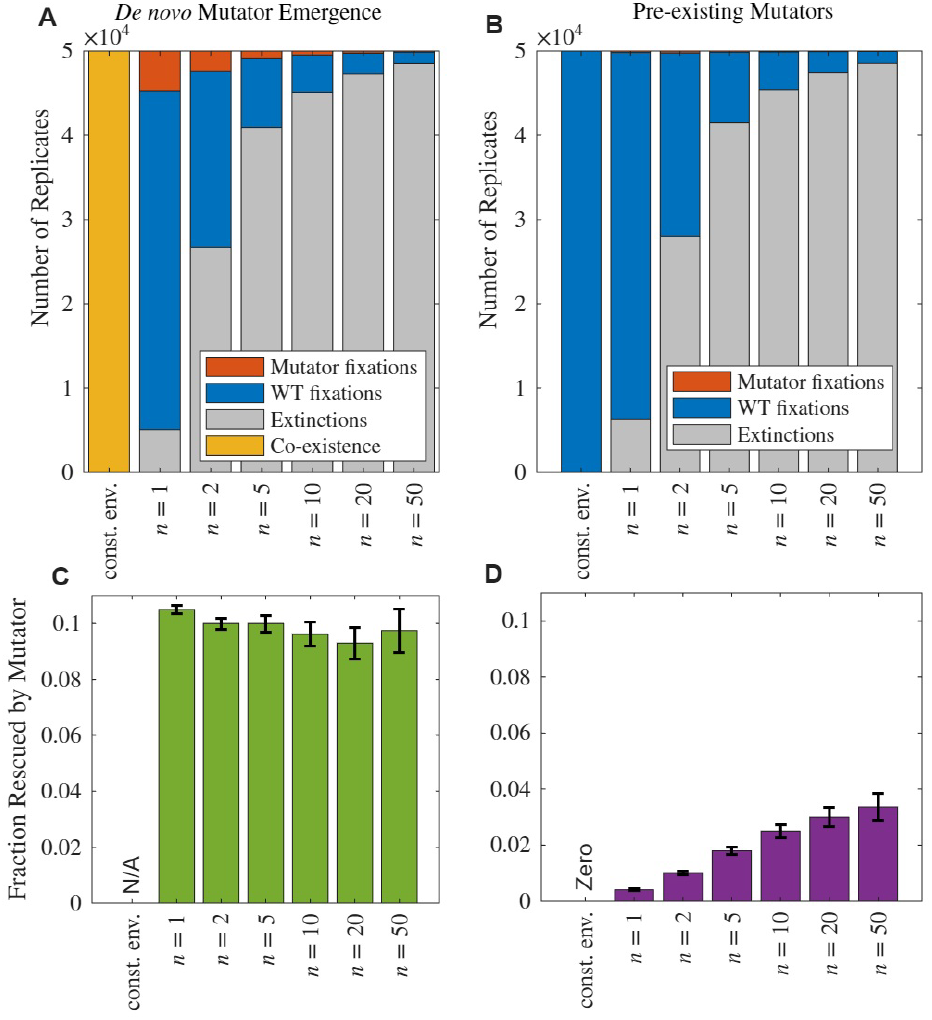
Environmental stress promotes mutator fixation through evolutionary rescue while no mutator fixations occur in constant environments (const. env.). (A, B) Stacked bar plots show outcomes of evolutionary rescue simulations under varying environmental change speeds (*n*), comparing (A) *de novo* mutator emergence and (B) pre-existing mutators. Mutator fixation (red) only occurs when environmental stress is present. (C, D) Fraction of populations rescued by mutators out of all rescued instances as a function of *n*, showing that pre-existing mutators increasingly contribute to rescue at higher *n* (D) whereas no such effect is seen for *de novo* mutators. Error bars represent standard error across 50,000 simulation replicates.

The fraction of populations rescued by *de novo* mutators, out of all instances in which the population is rescued (see Supplementary Information section S1.3), remains almost stable for fast or slow environmental changes (fig. 3C). Meanwhile, panel D shows that the fraction of pre-existing mutators contributing to rescue increases consistently with *n*, indicating that rapid environmental change amplifies the advantage of mutators already present in the population. This is due to the fact that faster environmental changes increase the growth rate of the rescue allele earlier, and pre-existing mutators have a higher fraction in the population the earlier this happens. For *de novo* emergence, however, the mutator fraction in the population stays at mutation-selection balance, as it emerges from the WT population at a constant rate. Trajectories of simulated populations through time for several illustrative cases are shown in Supplementary Section S1.3.1 and figures S1-S5.

### Mutator Emergence Increases Survival but is Limited by Mutation Costs

Figure 4A shows both analytical and simulation results for rescue probabilities versus the wildtype mutation rate. In the absence of mutators, the probability of rescuing the population (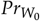, black line), increases superlinearly with the wildtype’s mutation rate until its growth rate starts to decline due to the cost of mutations, slowing the rise in the probability of rescue, and eventually leading to a decline in the chances of survival close to the critical mutation rate, *ν*_*c*_ (fig. 4A).

**Figure 4.**
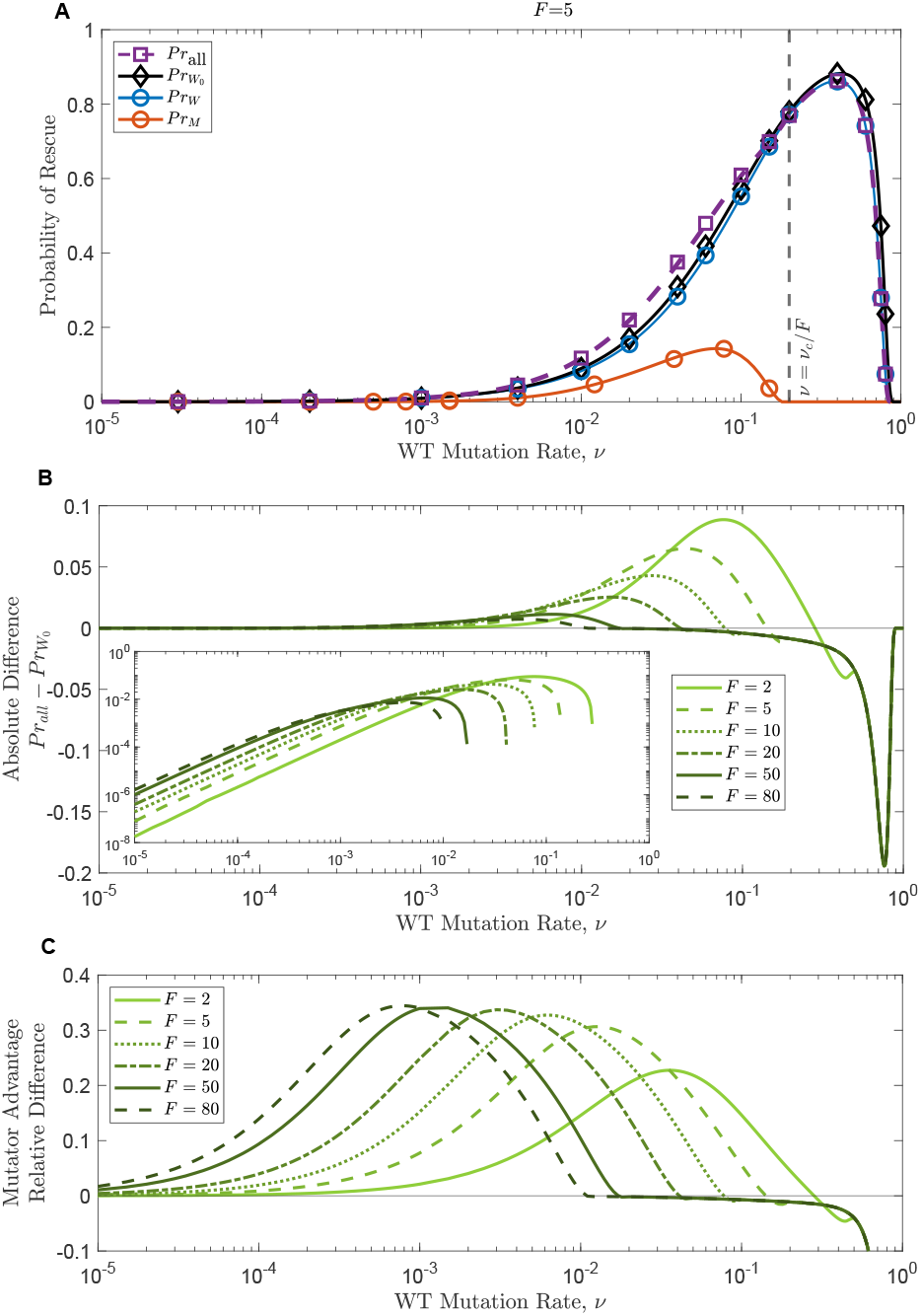
Allowing mutator emergence increases the overall survival chances at mild wildtype mutation rates. A: A mutator with strength *F* = 5 decreases the probability of rescue by the wildtype (blue vs black lines, 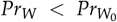), but increases the overall survival chances (purple, *Pr*_all_) as long as its growth rate is higher than its death rate (left of *ν* = *ν*_*c*_ /*F*), where it has some chance of rescuing the population (red, *Pr*_*M*_). B: The absolute difference between *Pr*_all_ and 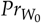. The inset shows the same plot on a loglog-scale. C: The relative advantage of allowing mutators (eq. 12) peaks at intermediate wildtype mutation rates as the strength of the mutator increases (darker green lines). Other parameters are *n* = 10, *E* = 2000, *θ* = 500 and death rates are 0.1. Symbols in panel A represent the simulation results for over 3 *×* 10^4^ replicates.

Introducing a mutator with strength *F* through *de novo* mutations leads to a slight decline in the probability of rescue by the wildtype, *Pr*_*W*_, due to the reduced wildtype population size (blue line lower than black line in fig. 4A). This also gives the mutator a chance to rescue the population as long as the wildtype’s mutation rate *ν* is less than *ν*_*c*_ /*F*, since the mutator has a growth rate of zero otherwise. Again, the closer the value of *ν* to *ν*_*c*_ /*F*, the more the increase in the probability of rescue by the mutator *Pr*_*M*_ is reduced, until it declines to zero once the mutator growth rate is less than or equal to the death rate (red line in fig. 4A).

Allowing mutators to emerge significantly boosts evolutionary rescue probabilities at intermediate wildtype mutation rates (fig. 4B,C). When mutation rates are initially low, survival probability are extremely low, but mutators enhance access to rescuing mutations (fig. 4B), increasing overall survival chances. The increase is modest in magnitude at these low mutation rates, but becomes relatively significant even in regions where survival seems very unlikely (e.g. around *ν* = 10^−4^ in fig. 4. On the other hand, this advantage diminishes as the wildtype mutation rate increases, due to growing mutation-related fitness costs.

### Stronger Mutators Thrive at Low Wildtype Mutation Rates, and Vice Versa

Mutation rates induce costs that can vary from being negligible at low mutation rates to significant at high mutation rates. Since the mutator’s mutation rate, *ν*^*′*^, is linked with the wildtype’s mutation rate such that *ν*^*′*^ = *Fν*, the cost for the mutator depends on both its strength, *F*, and the wildtype’s mutation rate, *ν*. As a result, increasing the mutator strength promotes rescue by mutators at low wildtype mutation rates, but this changes as *ν* increases leading eventually to an opposite effect, where higher mutator strengths become more detrimental due to their significant costs (fig. 5).

**Figure 5.**
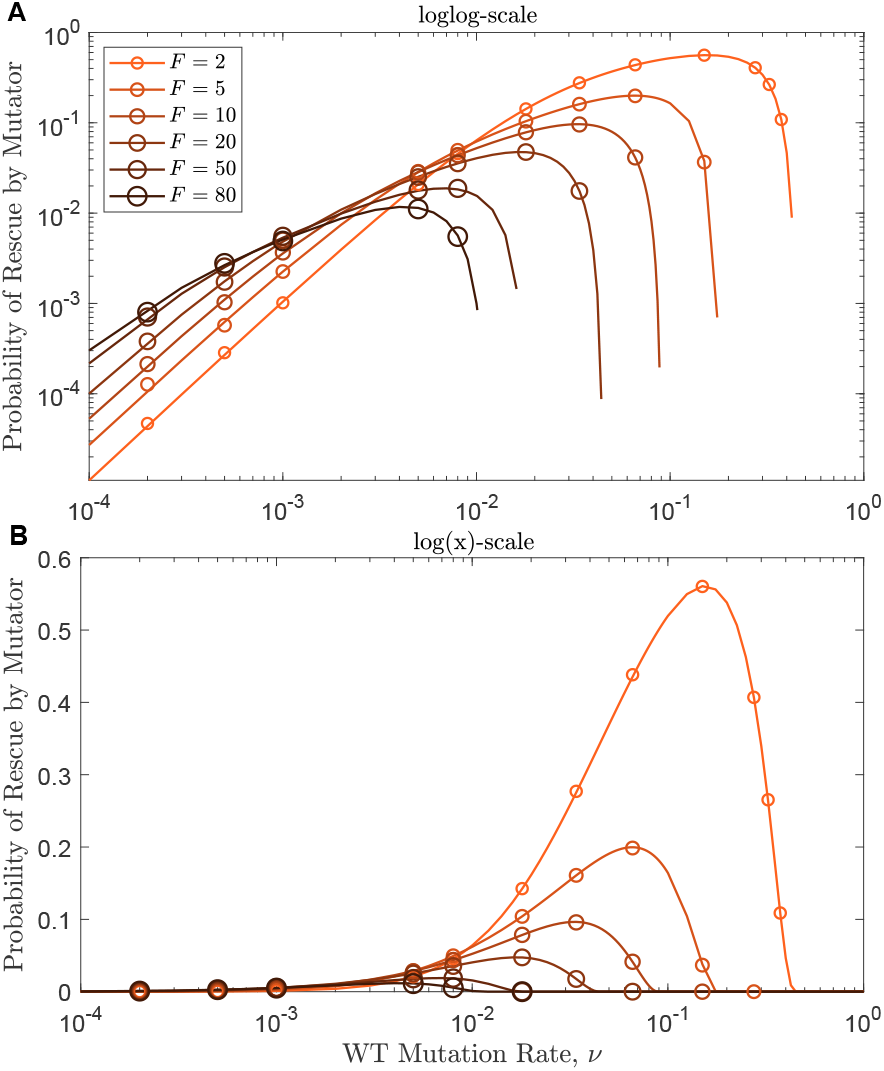
Elevating mutation rates from low wild-type levels incurs minimal costs, thus favouring larger mutation rate multipliers, while this effect flips for high wildtype mutation rates. Single mutators with varying strengths emerge with an emergence factor *E* = 500. Other parameters are: *n* = 2, *θ* = 500 and death rates are 0.1. Results are shown on a loglog-scale in A and on a log(x)-scale in B. Circles show simulation results of over 15 *×* 10^4^.

This is true when single mutator populations confer rescue (fig. 5), where the value of *F* determines the flipping point. It also applies when multiple mutator populations are introduced (see Supplement section S2).

### Lower Mutator Emergence Rates Reduce Probability of Rescue by the Mutator

On the other hand, varying a parameter that only affects the mutator population does not change the probability of rescue by the wildtype much. We see this when varying the mutator emergence rates by changing the value of *e* (fig. 6A), or by varying the mutation rate multiple *F* (not shown; very similar to fig. 6A). In contrast, varying the mutator emergence rate has a large effect on the probability of rescue by the mutator, increasing from very low probabilities for low values of *e* and reaching almost 0.6 for the highest value *e* = 2000 for the parameters of choice in fig. 6B. Hence, as one would expect, the mutator advantage increases for higher mutator emergence rates, except (of course) when the cost of the mutator’s mutation rate reduces its growth rate significantly (fig. 6C).

**Figure 6.**
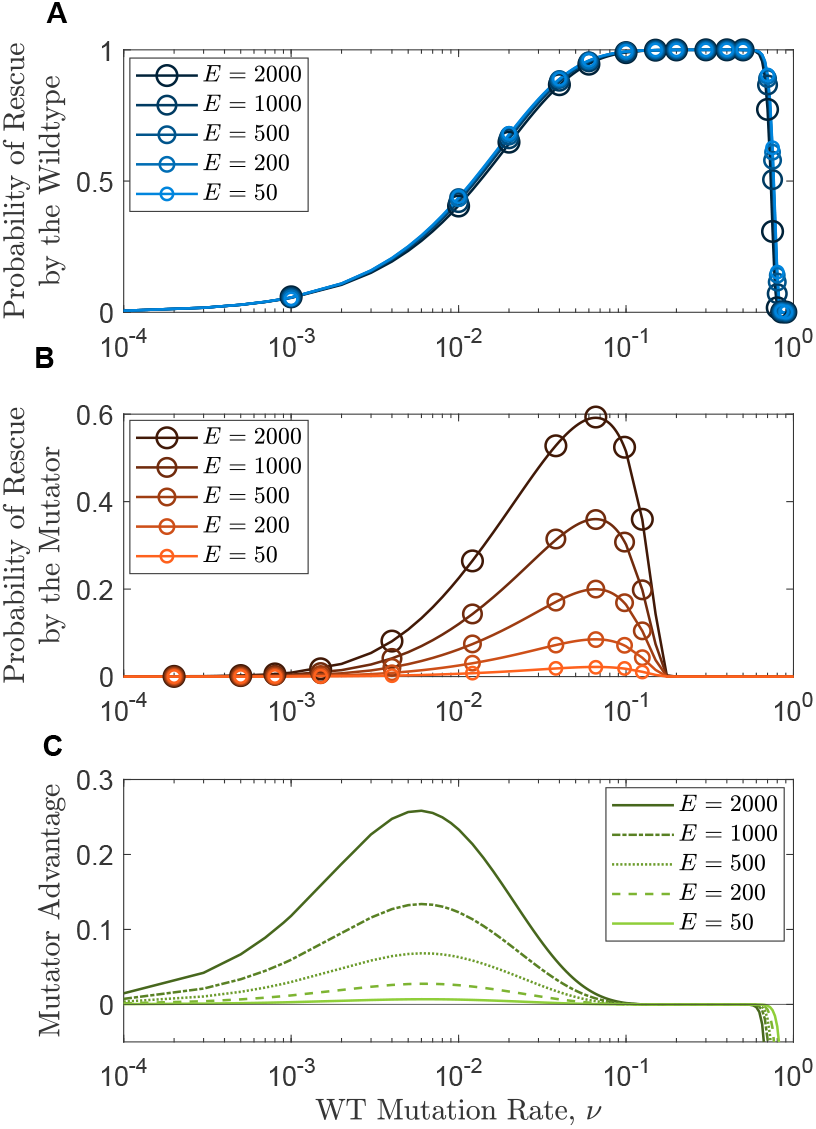
Lower mutator emergence rates reduce probability of rescue by the mutator. A,B: Higher mutator emergence rates show no significant effect on the probability of rescue by the wildtype, while notably elevating rescue by mutators. C: Overall, increased emergence rates are beneficial for the population survival chances as long as mutators do not have zero growth rates. Other parameters are: *n* = 2, *F* = 5, *θ* = 500 and death rates are 0.1. Circles show simulation results for over 10^4^ replicates for wildtype and over 25 *×* 10^4^ for the mutator.

### The Pace of Environmental Change Modulates Mutator Benefits

For rapid environmental changes (high *n* values), rescue mutants start to have positive growth rates closer to the centered time of environmental change, *θ*, and only then they can grow (fig. 1). Thus, the faster the environmental change, the lower the chances that rescue mutant lineages occur and grow, generally leading to lower probabilities of rescue by either the wildtype or the mutator (fig. 7A,B).

**Figure 7.**
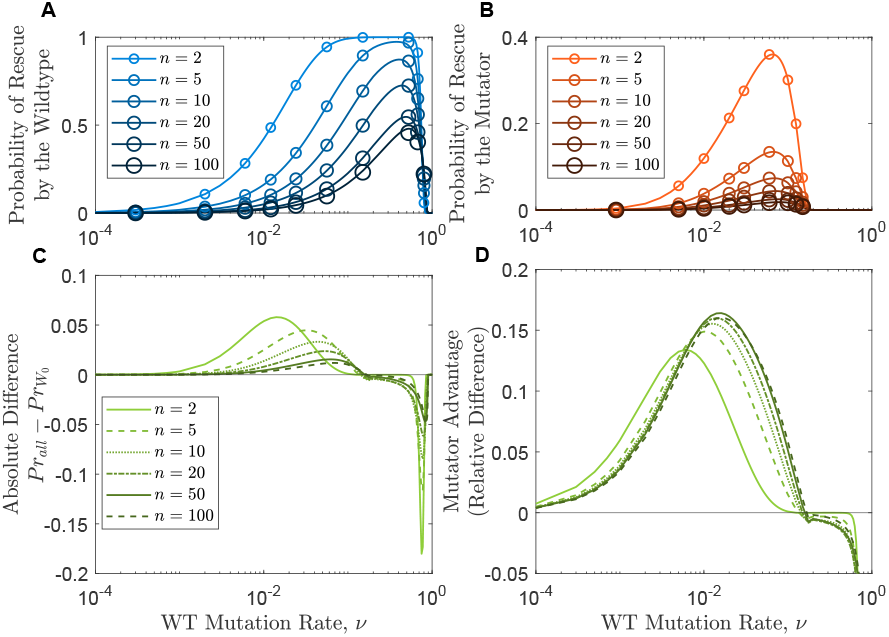
More sudden environmental changes reduce the probabilities of rescue, but confer larger advantages at intermediate wildtype mutation rates. A,B: Probability of rescue by the wildtype or by the mutator decreases significantly for faster environmental changes. C,D: more gradual environmental changes promote mutators more at low wildtype mutation rates, and vice versa. Other parameters are: *E* = 1000, *F* = 5, *θ* = 500 and death rates are 0.1. Circles show simulation results for over 10^4^ replicates for wildtype and over 25 *×* 10^4^ for the mutator.

Counterintuitively, the advantage of introducing mutators is not always higher for more sudden environmental changes (fig. 7C,D). At low wildtype mutation rates, mutator emergence is more advantageous when the environment changes more slowly, while the opposite is true for higher wildtype mutation rates, unless it is too high, flipping the advantage to a disadvantage.

Taking a closer look at how this advantage changes between gradual or abrupt environmental changes (fig. 8), we see a region of advantageous mutator emergence (bright green) at low

**Figure 8.**
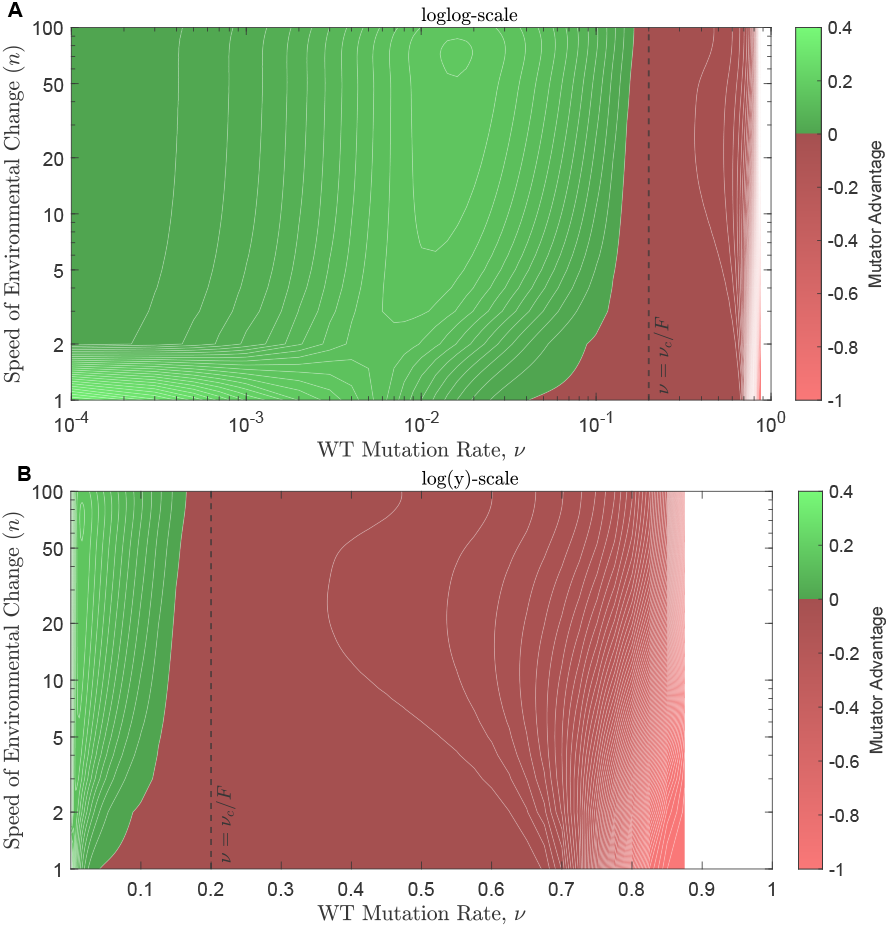
Highly advantageous mutator emergence only occurs at low wildtype mutation rates when the environment changes very slowly, while milder advantages take place at intermediate wildtype mutation rates no matter how fast the environment changes. The same region is shown in A and B, on a log scale for both axes in A, but on only the *y*-axis in B. Parameters are: *F* = 5, *θ* = 500, *E* = 1000, and death rates are 0.1.

wildtype mutation rates and very slow environmental changes (*n <* 2), and this advantage increases sharply as *n* gets even smaller. As wildtype mutation rates increase to intermediate levels, mutators start contributing more to survival at faster environmental changes. This advantage starts to fade again as the mutation rates get closer to the critical value, *ν* = *ν*_*c*_ /*F*, at which point mutators no longer survive. Introducing mutators becomes increasingly disadvantageous after that, hindering survival more at slower environmental changes. The mutator advantage becomes undefined when the probability of rescue by the wildtype in the absence of mutators, 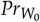, drops to zero.

A sensitivity analysis for the choice of the inflection point *θ* in the Hill function for the growth rate is presented in supplementary section S1.3.2. Since populations do not start at equilibrium (see fig. S1), the choice of *θ* can affect the outcome if the environmental change starts before populations reach steady state.

### Strong Pre-existing Mutators have Significant Advantages at Low Wildtype Mutation Rates

Figure 9 shows that when mutators exist before the environmental change, they can substantially increase rescue probabilities. Pre-existing mutators contribute more to rescue when the environmental change occurs earlier (fig. 9A,B), presumably due to the decline in the mutator population over time.

**Figure 9.**
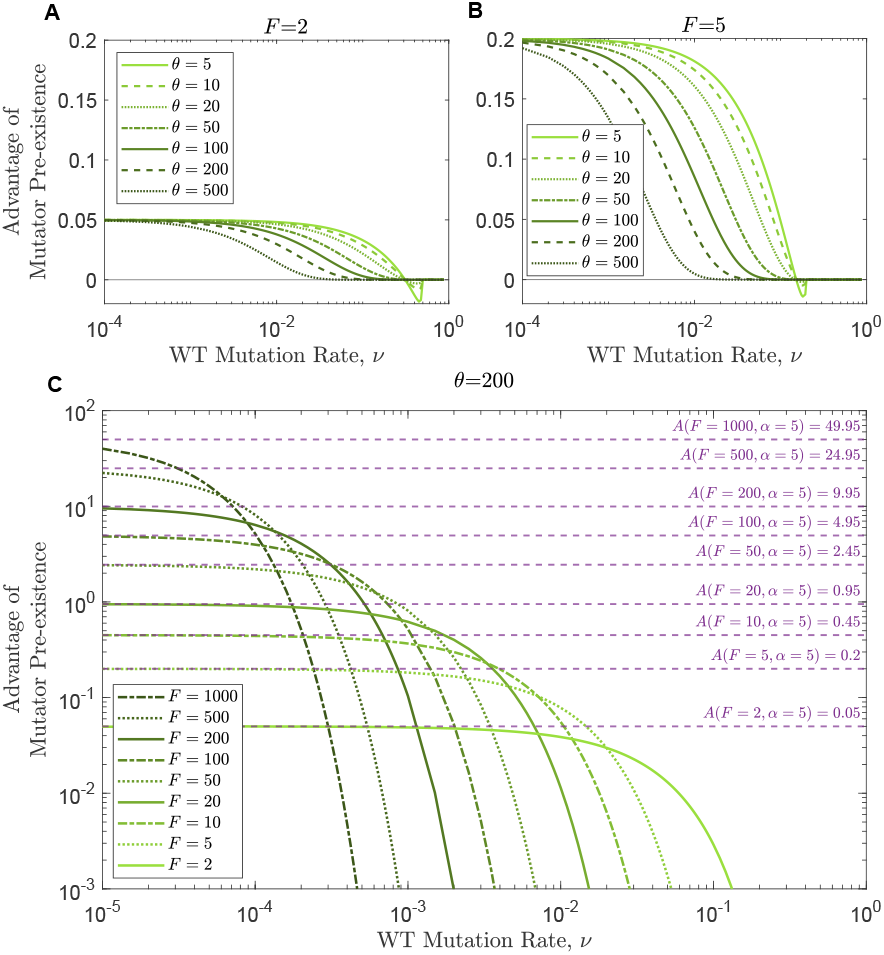
Strong pre-existing mutators have significant advantages at low wildtype mutation rates. A,B: The advantage of mutator pre-existence (with strength *F* = 2 and *F* = 5, respectively) is higher for earlier environmental changes, and reaches its maximum at low wildtype mutation rates as it converges to *A*(*F, α*) (eq. 21). Other parameters are *n* = 2 and death rates are 0.1. C: Mutator pre-existence provides a significant advantage at low wildtype mutation rates, which converges to *A*(*F, α*) (eq. 21), but higher wildtype mutation rates gradually eradicate this effect due to the cost of mutation. Parameters are: *n* = 10, *θ* = 200, and death rates are 0.1

Moreover, the wave effect (local maximum in *ν*) that is seen in the advantage of emerging mutators (fig. 4) disappears when mutators pre-exist. Instead, this advantage approaches an upper limit at low wildtype mutation rates, which is expected since lower wildtype mutation rates no longer reduce the mutator population when mutators pre-exist. This limit is higher when the mutator’s strength is higher (fig. 9C), and closely agrees with the mutator advantage predicted in eq. (21). We note that that prediction assumes no cost of mutations, an assumption that is most closely realized when *ν* is low (fig. 9C).

## Discussion

Mutation rates are known to play a critical role in evolutionary rescue (Orr and Unckless 2008; Tanaka and Wahl 2022; Marrec and Bitbol 2020a; Anciaux et al. 2019). In general, higher mutation rates increase access to rescuing mutants, and thus increase the probability of rescue (Marrec and Bitbol 2020a), as long as they do not reach a level leading to lethal mutagenesis or mutational meltdown (Anciaux et al. 2019). On the other hand, the contribution of mutators to adaptation has been well studied (Taddei *et al*. 1997a; Travis and Travis 2002; Raynes *et al*. 2011), typically confirming that mutators facilitate adaptation when sufficient beneficial mutations are available (Ram and Hadany 2012; Tuffaha *et al*. 2023), but that they are selected against once the environment stabilizes (Kessler and Levine 1998). Nevertheless, little work has investigated how having mutators within a population affects evolutionary rescue probabilities, or, asked differently, how severe environmental shifts promote mutator emergence. By providing a theoretical framework for predicting mutator emergence, our study offers insights into optimizing treatment strategies that minimize unintended mutator promotion.

Taken together, our results highlight that mutators are unlikely to persist at high frequencies in a stable environment due to mutational costs, but can be promoted under environmental stress through evolutionary rescue events. This finding underscores the critical role of selective pressures in driving the emergence and fixation of high-mutation-rate lineages. Thus, **evolutionary rescue promotes mutators**.

Conversely, we have shown that **mutators promote evolutionary rescue**. Allowing the emergence of mutators can substantially increase the overall probability of rescue (up to 40% in the parameter regime we investigated, fig. 4B), but the conditions under which this benefit can be reached depend on many factors. As one would expect, the higher the mutator emergence rate the higher the advantage of mutators (fig. 6C), but stronger emerging mutators (those with higher mutation rates) are not always more beneficial due to mutation rate costs (fig. 4B). In fact, stronger mutators are more likely to rescue the population, and thus emerge in response to environmental stress, if the wildtype mutation rate is quite low, while this effect flips for wildtype strains with higher mutation rates (fig. 5).

Our model imposes the cost of mutation as a reduction in growth rate, which can be caused by increased deleterious load, as well as the burden of lethal mutations. One more risk caused by high mutation rates is the increased rate of back mutations at the rescue allele, which would be an extra burden of highly deleterious, or even lethal, mutations available only to rescuing mutants. Thus in reality we do not expect the rescuing mutants from mutator backgrounds to reach exactly the same maximum growth rates as their mutator ancestors. Although in this study we have assumed the same maximum fitness can be reached, each in its favored environment, we do not expect this assumption to lead to qualitative changes in the results. If back mutations have the same rate as mutations to the rescue allele (they can have different rates (Tanaka and Wahl 2022)), back mutations on rescuing mutator backgrounds would only occur at rate *Frν*, so they should be as rare as rescue mutations and are not expected to measurably affect fitness.

In our model, the mutation rate of mutator lineages is defined as a constant multiple of the wildtype mutation rate, *ν*^*′*^ = *Fν*. This multiplicative formulation is both biologically motivated and standard in the theoretical literature. Empirically, mutator phenotypes commonly arise from disruptions in genome-wide error correction mechanisms—such as loss-of-function mutations in DNA proofreading or mismatch repair genes (e.g., *mutS, mutL, dnaQ*)—and these lead to proportional increases in mutation rates across the genome (Sane *et al*. 2024; Boyce 2022; Chopra *et al*. 2003). Consequently, theoretical studies modeling the evolution or impact of mutators routinely adopt this multiplicative relationship (Lynch 2011; Anciaux *et al*. 2019; Wylie *et al*. 2009b).

We note, however, that a potential drawback of the multiplicative formulation is that it amplifies the difference between the wildtype and mutator mutation rates as *ν* increases. This means the mutator advantage may appear stronger at moderate to high wildtype mutation rates by construction, even in the absence of mechanistic changes. Conversely, the difference is attenuated when *ν* is low. While this behavior is consistent with biological expectations under most known mutator mechanisms, it is important to interpret the trends involving variation in *ν* with this structural dependency in mind. That said, other relationships between the wildtype and mutator mutation rates may arise in nature. For instance, under a modular repair architecture in which each repair pathway is responsible for correcting a fixed fraction of replication errors (Peterson and Almouzni 2013; Uphoff and Kapanidis 2014; Luo et al. 2010), the loss of a specific repair pathway could lead to an additive increase in mutation rate—i.e., a fixed increment regardless of baseline fidelity. Importantly, our modeling framework is flexible and can readily accommodate alternative formulations such as this. To illustrate, we provide an example in Supplementary Section S1.5 using an additive mutator model where *ν*^*′*^ = *ν* + Δ, and observe how the results compare to the multiplicative case.

The shape and timing of environmental deterioration also play a critical role in shaping evolutionary dynamics. In our framework, we model environmental change using a Hill function, where the coefficient *n* controls the steepness but also affects the onset of decline. Higher values of *n* lead to more abrupt but delayed changes, creating scenarios in which “rapidly deteriorating” environments remain benign for longer than gradually changing ones (fig 1). This temporal asymmetry may help explain some of the non-monotonic patterns observed in mutator advantage (e.g., fig. 7 and 8). Additionally, in Supplementary Figures S6, S7, we explore the behavior of rescue probabilities under early environmental changes versus long-delay regimes (i.e., smaller versus larger *θ*). In the pre-existing case, mutator lineages decline over time due to selection, with rare stochastic persistence, resulting in a slow convergence of rescue probabilities to zero. In contrast, in the *de novo* case, the absence of back mutation allows rescue lineages to accumulate during extended time intervals, leading to a steady increase in rescue probability. These results highlight that both the form of environmental change and assumptions about mutation reversibility can substantially influence long-term outcomes, and merit further exploration in future models.

The balance between the costs and benefits of elevated mutation rates is critically important in medical contexts such as cancer evolution and antimicrobial therapy. In both cases, therapeutic agents are sometimes designed or observed to increase mutation rates—either as a byproduct of DNA-damaging effects or as a deliberate strategy to accelerate mutational meltdown. However, this approach is double-edged. While high mutation rates can indeed be lethal due to mutational overload or error catastrophe (Anciaux et al. 2019), they can also inadvertently promote *evolutionary rescue*, enabling populations to survive otherwise lethal treatments. Recent work by Kuosmanen et al. (Kuosmanen et al. 2021) highlights this trade-off: they show that drug regimens inducing high mutagenesis can paradoxically increase the likelihood of resistance emergence. Their findings emphasize that intermediate dosing may offer the best balance—maximizing treatment efficacy while minimizing the chances of escape through de novo mutations. Our results add to this discussion by examining the role of pre-existing mutator alleles, showing that the sudden appearance or presence of high mutation rate variants can substantially raise the chances of evolutionary rescue—particularly when the wildtype mutation rate is low. These findings suggest that therapeutic strategies must carefully quantify the levels of mutagenicity they induce, and take into account not only the average mutation rate, but also the potential for standing variation in mutation rate modifiers within a population. Inadvertently selecting for or enabling mutators may shift the evolutionary balance in favor of escape, undermining the long-term effectiveness of treatment. Thus, our work reinforces the need for quantitative evolutionary models when designing therapies that interact with mutational processes.

For the reasons outlined above, it is common for a mutator “advantage” to be seen as an increased risk introduced by mutators in the resistance context. Greenspoon and Mideo (2017) explored this risk in the framework of drug resistance evolution. Although they explore different levels of fitness costs of mutator alleles, they keep the cost fixed and do not assume, as we do, that the cost increases for stronger mutators. According to their simulations, mutator emergence poses higher risks for low recombination rates, high pathogen virulence, low environmental fluctuation rates, and when re-infection by different parasite strains is more common, which are factors not considered in our model. Similar to their findings (Greenspoon and Mideo 2017), we show that in the “probability of evolutionary rescue” space, mutator evolution appears to be significant only in the intermediate regions where the likelihood is neither very high (where rescue is already almost guaranteed by the wildtype) nor very low (fig. 4, fig. 6, fig 7). As Greenspoon and Mideo suggest, this finding can potentially explain the lack of mutator advantage sometimes observed in clinical settings (Auerbach *et al*. 2015), while this advantage is very clear in many experimental studies (Eliopoulos and Blázquez 2003; Giraud *et al*. 2002; Gifford *et al*. 2023).

Our work shows that mutator pre-existence, unlike mutator *de novo* emergence, can significantly promote survival probabilities, especially when these probabilities are small in the absence of mutators (fig 9). This can partially explain the high number of rescue events by Δ*mutS* mutators (*F* = 80) that were introduced by Gifford et. al. (2023) at 5-30% frequencies to test their potential in promoting single- or double-resistance alleles that can withstand two different antibiotics. Mutators in that study sampled more specialized resistance mutations compared to the wildtype, and showed further elevations in mutation rate in the rescuing population, which was not seen in the wildtype background. This additional advantage of mutators’ increased access to specialized rescue mutations and also to other mutation-rate-modifier loci is not captured by our model. If mutators pre-existed and also emerged in the wildtype or mutator backgrounds, this is expected to potentially confer further mutator advantage at both low and intermediate wildtype mutation rates, and a mathematical model for this is worth developing in future work.

Although the pronounced advantage of pre-existing mutators in our results mirrors the experimental findings of Gifford *et al*. (2023), our *de novo* analysis cautions that extending those laboratory insights to natural populations must be done with care. In nature, mutators (especially strong ones) are only expected to co-exist with the wildtype at very low frequencies in constant environments (Boe *et al*. 2000; Taddei *et al*. 1997b; Miller 1996). Thus, if mutators are introduced at higher frequencies without an environmental change, they are expected to be purged by selection until they reach mutation-selection balance again (Desai and Fisher 2011; Sprouffske et al. 2018). In drugresistance-evolution experiments, such as Gifford *et al*. (2023), mutators likely have higher chances of rescue compared to natural environments due to their introduction at high frequencies, and possibly also due to the relatively large fraction of beneficial mutations available.

In fact, Gifford *et al*. (2023) found that mutator strains developed antibiotic resistance by more specialized mutations compared to the wildtype, which showed more generalist rescue mutants. This can be partially explained by potential mutational spectra shifts in these mutators that could have enhanced their access to these specialized mutations (Tuffaha et al. 2023). However, it has been shown that the Δ*mutS* strain that Gifford *et al*. (2023) introduced in their experiments reinforces the transition bias in its *E. coli* K12 ancestor (Sane et al. 2024). On the other hand, Gifford *et al*. (2023) reported further elevations in mutation rates that occurred in rescuing mutants from this mutator’s background, which could have potentially been coupled with bias reversals. Investigating this more complex effect of sequential elevations in mutation rates and bias shifts is an interesting research avenue.

An increased mutator advantage corresponds to a higher ratio of instances in which a population is rescued by the mutator to the instances it is rescued by the wild type. Thus a higher mutator advantage reflects greater chances for mutator emergence in response to environmental stress. These chances are the highest when the environmental change is very slow (*n* = 1, fig. 8A) and at low wildtype mutation rates, but these slow environmental changes also result in the strongest disadvantage for mutators when the wildtype itself has a high mutation rate (*n* = 1, fig. 8B).

As the environment starts changing faster, the mutator advantage is rapidly reduced at low mutation rates (fig. 8A), presumably due to the two mutational steps needed for the mutator to rescue the population: the mutator needs to emerge first and then mutate at the rescue allele, which are two sequential rare events in that regime. Yet, intermediate wildtype mutation rates give this two-step mutational process higher chances to occur at fast environmental changes, allowing such sudden changes to promote mutator emergence.

In reality, at very high mutation rates, anti-mutator alleles (Bak et al. 2014; Martincorena and Luscombe 2013) might be favored, leading to lower mutation rates, reducing mutation cost, and probably promoting survival. In that sense, harsh environmental stresses on mutators would be likely to promote anti-mutator phenotypes, an insight worth investigating in future work.

## Code Availability

Simulation codes can be found in https://github.com/MarwaTuffaha/MutatorRescue.

## Literature cited

Alejandre C, Calle-Espinosa J, Iranzo J. 2024. Synergistic epistasis among cancer drivers can rescue early tumors from the accumulation of deleterious passengers. PLOS Computational Biology. 20:e1012081.

Alexander H, Martin G, Martin O, Bonhoeffer S. 2014. Evolutionary rescue: linking theory for conservation and medicine. evol. appl. 7: 1161–1179.

Anciaux Y, Lambert A, Ronce O, Roques L, Martin G. 2019. Population persistence under high mutation rate: from evolutionary rescue to lethal mutagenesis. Evolution. 73:1517–1532.

Auerbach A, Kerem E, Assous MV, Picard E, Bar-Meir M. 2015. Is infection with hypermutable pseudomonas aeruginosa clinically significant? Journal of Cystic Fibrosis. 14:347–352.

Bak ST, Sakellariou D, Pena-Diaz J. 2014. The dual nature of mismatch repair as antimutator and mutator: for better or for worse. Frontiers in genetics. 5:287.

Baquero F, Martinez JL, F. Lanza V, Rodríguez-Beltrán J, Galán JC, San Millán A, Cantón R, Coque TM. 2021. Evolutionary pathways and trajectories in antibiotic resistance. Clinical Microbiology Reviews. 34:e00050–19.

Bell G. 2017. Evolutionary rescue. Annual Review of Ecology, Evolution, and Systematics. 48:605–627.

Blattner FR, Plunkett III G, Bloch CA, Perna NT, Burland V, Riley M, Collado-Vides J, Glasner JD, Rode CK, Mayhew GF et al. 1997. The complete genome sequence of escherichia coli k-12. science. 277:1453–1462.

Boe L, Danielsen M, Knudsen S, Petersen JB, Maymann J, Jensen PR. 2000. The frequency of mutators in populations of escherichia coli. Mutation Research/Fundamental and Molecular Mechanisms of Mutagenesis. 448:47–55.

Boyce KJ. 2022. Mutators enhance adaptive micro-evolution in pathogenic microbes. Microorganisms. 10:442.

Callaghan TV, Bergholm F, Christensen TR, Jonasson C, Kokfelt U, Johansson M. 2010. A new climate era in the sub-arctic: Accelerating climate changes and multiple impacts. Geophysical Research Letters. 37.

Carlson SM, Cunningham CJ, Westley PA. 2014. Evolutionary rescue in a changing world. Trends in ecology & evolution. 29:521–530.

Chevin LM, Lande R, Mace GM. 2010. Adaptation, plasticity, and extinction in a changing environment: towards a predictive theory. PLoS biology. 8:e1000357.

Chopra I, O’Neill AJ, Miller K. 2003. The role of mutators in the emergence of antibiotic-resistant bacteria. Drug Resistance Updates. 6:137–145.

Couce A, Guelfo JR, Blázquez J. 2013. Mutational spectrum drives the rise of mutator bacteria. PLoS Genetics. 9:e1003167.

DeFilippo LB, McManus LC, Schindler DE, Pinsky ML, Colton MA, Fox HE, Tekwa E, Palumbi SR, Essington TE, Webster MM. 2022. Assessing the potential for demographic restoration and assisted evolution to build climate resilience in coral reefs. Ecological applications. 32:e2650.

Denamur E, Matic I. 2006. Evolution of mutation rates in bacteria. Molecular microbiology. 60:820–827.

Desai MM, Fisher DS. 2011. The balance between mutators and nonmutators in asexual populations. Genetics. 188:997–1014.

Eliopoulos GM, Blázquez J. 2003. Hypermutation as a factor contributing to the acquisition of antimicrobial resistance. Clinical Infectious Diseases. 37:1201–1209.

Feiner N, Brun-Usan M, Uller T. 2021. Evolvability and evolutionary rescue. Evolution & Development. 23:308–319.

Ferrare JT, Good BH. 2024. Evolution of evolvability in rapidly adapting populations. Nature Ecology & Evolution. pp. 1–12.

Garnier J, Cotto O, Bouin E, Bourgeron T, Lepoutre T, Ronce O, Calvez V. 2023. Adaptation of a quantitative trait to a changing environment: new analytical insights on the asexual and infinitesimal sexual models. Theoretical Population Biology. 152:1–22.

Gerrish PJ, García-Lerma JG. 2003. Mutation rate and the efficacy of antimicrobial drug treatment. The Lancet infectious diseases. 3:28–32.

Gifford DR, Berríos-Caro E, Joerres C, Suñé M, Forsyth JH, Bhattacharyya A, Galla T, Knight CG. 2023. Mutators can drive the evolution of multi-resistance to antibiotics. PLoS Genetics. 19:e1010791.

Gillespie DT. 1977. Exact stochastic simulation of coupled chemical reactions. The journal of physical chemistry. 81:2340–2361.

Giraud A, Matic I, Radman M, Fons M, Taddei F. 2002. Mutator bacteria as a risk factor in treatment of infectious diseases. Antimicrobial agents and chemotherapy. 46:863–865.

Gomulkiewicz R, Holt RD. 1995. When does evolution by natural selection prevent extinction? Evolution. pp. 201–207.

Gonzalez A, Ronce O, Ferriere R, Hochberg ME. 2013. Evolutionary rescue: an emerging focus at the intersection between ecology and evolution.

Greenspoon PB, Mideo N. 2017. Evolutionary rescue of a parasite population by mutation rate evolution. Theoretical Population Biology. 117:64–75.

Greenspoon PB, Spencer HG. 2021. Avoiding extinction under nonlinear environmental change: models of evolutionary rescue with plasticity. Biology Letters. 17:20210459.

Hinsch M, Robertson DL, Silverman E. 2023. Evolutionary rescue effect disappears under more realistic assumptions. medRxiv. pp. 2023–05.

Holt RD, Gomulkiewicz R. 1997. The evolution of species’ niches: a population dynamic perspective. Case studies in mathematical modelling: ecology, physiology, and cell biology. pp. 25–50.

Jee J, Rasouly A, Shamovsky I, Akivis Y, R. Steinman S, Mishra B, Nudler E. 2016. Rates and mechanisms of bacterial mutagenesis from maximum-depth sequencing. Nature. 534:693–696.

Jiao J, Gilchrist MA, Fefferman NH. 2020. The impact of host metapopulation structure on short-term evolutionary rescue in the face of a novel pathogenic threat. Global Ecology and Conservation. 23:e01174.

Kessler DA, Levine H. 1998. Mutator dynamics on a smooth evolutionary landscape. Physical review letters. 80:2012.

Kuosmanen T, Cairns J, Noble R, Beerenwinkel N, Mononen T, Mustonen V. 2021. Drug-induced resistance evolution necessitates less aggressive treatment. PLoS computational biology. 17:e1009418.

Lukjancenko O, Wassenaar TM, Ussery DW. 2010. Comparison of 61 sequenced escherichia coli genomes. Microbial ecology. 60:708–720.

Luo M, He H, Kelley MR, Georgiadis MM. 2010. Redox regulation of dna repair: implications for human health and cancer therapeutic development. Antioxidants & redox signaling. 12:1247–1269.

Lynch M. 2011. The lower bound to the evolution of mutation rates. Genome Biology and Evolution. 3:1107–1118.

Maciá MD, Blanquer D, Togores B, Sauleda J, Pérez JL, Oliver A. 2005. Hypermutation is a key factor in development of multiple-antimicrobial resistance in pseudomonas aeruginosa strains causing chronic lung infections. Antimicrobial agents and chemotherapy. 49:3382–3386.

Marrec L, Bank C. 2023. Evolutionary rescue in a fluctuating environment: periodic versus quasi-periodic environmental changes. Proceedings of the Royal Society B: Biological Sciences. 290:20230770. Epub 2023 May 31.

Marrec L, Bitbol AF. 2020a. Adapt or perish: Evolutionary rescue in a gradually deteriorating environment. Genetics. 216:573–583.

Marrec L, Bitbol AF. 2020b. Resist or perish: fate of a microbial population subjected to a periodic presence of antimicrobial. PLoS computational biology. 16:e1007798.

Martincorena I, Luscombe NM. 2013. Non-random mutation: The evolution of targeted hypermutation and hypomutation. BioEssays. 35:123–130.

Martinez J, Baquero F. 2000. Mutation frequencies and antibiotic resistance. Antimicrobial agents and chemotherapy. 44:1771–1777.

Miller JH. 1996. Spontaneous mutators in bacteria: insights into pathways of mutagenesis and repair. Annual review of microbiology. 50:625–643.

Nerem RS, Beckley BD, Fasullo JT, Hamlington BD, Masters D, Mitchum GT. 2018. Climate-change–driven accelerated sealevel rise detected in the altimeter era. Proceedings of the national academy of sciences. 115:2022–2025.

Orive ME, Holt RD, Barfield M. 2019. Evolutionary rescue in a linearly changing environment: limits on predictability. Bulletin of Mathematical Biology. 81:4821–4839.

Orr HA, Unckless RL. 2008. Population extinction and the genetics of adaptation. Amer. Nat.. 172:160–169.

Orr HA, Unckless RL. 2014. The population genetics of evolutionary rescue. PLoS Genetics. 10.

Osmond MM, Otto SP, Martin G. 2020. Genetic paths to evolutionary rescue and the distribution of fitness effects along them. Genetics. 214:493–510.

O’Dea RE, Noble DW, Johnson SL, Hesselson D, Nakagawa S. 2016. The role of non-genetic inheritance in evolutionary rescue: epigenetic buffering, heritable bet hedging and epigenetic traps. Environmental epigenetics. 2:dvv014.

Peterson CL, Almouzni G. 2013. Nucleosome dynamics as modular systems that integrate dna damage and repair. Cold Spring Harbor perspectives in biology. 5:a012658.

Ram Y, Hadany L. 2012. The evolution of stress-induced hypermutation in asexual populations. Evolution. 66:2315–2328.

Ramiro RS, Durão P, Bank C, Gordo I. 2020. Low mutational load and high mutation rate variation in gut commensal bacteria. PLoS biology. 18:e3000617.

Raynes Y, Gazzara MR, Sniegowski PD. 2011. Mutator dynamics in sexual and asexual experimental populations of yeast. BMC evolutionary biology. 11:1–7.

Raynes Y, Sniegowski P. 2014. Experimental evolution and the dynamics of genomic mutation rate modifiers. Heredity. 113:375–380.

Romero-Mujalli D, Jeltsch F, Tiedemann R. 2019. Elevated mutation rates are unlikely to evolve in sexual species, not even under rapid environmental change. BMC Evolutionary Biology. 19:1–9.

Sane M, Diwan GD, Bhat BA, Wahl LM, Agashe D. 2023. Shifts in mutation spectra enhance access to beneficial mutations. Proceedings of the National Academy of Sciences..

Sane M, Parveen S, Agashe D. 2024. Mutation bias alters the distribution of fitness effects of mutations. bioRxiv. pp. 2024–03.

Evolutionary Rescue Promotes Mutators Shaver AC, Dombrowski PG, Sweeney JY, Treis T, Zappala RM, Sniegowski PD. 2002. Fitness evolution and the rise of mutator alleles in experimental escherichia coli populations. Genetics. 162:557–566.

Sniegowski PD, Gerrish PJ, Johnson T, Shaver A. 2000. The evolution of mutation rates: separating causes from consequences. Bioessays. 22:1057–1066.

Sniegowski PD, Gerrish PJ, Lenski RE. 1997. Evolution of high mutation rates in experimental populations of e. coli. Nature. 387:703–705.

Sprouffske K, Aguilar-Rodriguez J, Sniegowski P, Wagner A. 2018. High mutation rates limit evolutionary adaptation in escherichia coli. PLoS genetics. 14:e1007324.

Taddei F, Radman M, Maynard-Smith J, Toupance B, Gouyon PH, Godelle B. 1997a. Role of mutator alleles in adaptive evolution. Nature. 387:700–702.

Taddei F, Radman M, Maynard Smith J, Toupance B, Gouyon PH, Godelle B. 1997b. Role of mutator alleles in adaptive evolution. Nature. 387:700–702.

Tanaka MM, Wahl LM. 2022. Surviving environmental change: when increasing population size can increase extinction risk. Proceedings of the Royal Society B. 289:20220439.

Thompson DA, Desai MM, Murray AW. 2006. Ploidy controls the success of mutators and nature of mutations during budding yeast evolution. Current Biology. 16:1581–1590.

Tomasini M, Peischl S. 2022. The role of spatial structure in multideme models of evolutionary rescue. Journal of evolutionary biology. 35:986–1001.

Travis J, Travis E. 2002. Mutator dynamics in fluctuating environments. Proceedings of the Royal Society of London. Series B: Biological Sciences. 269:591–597.

Tuffaha MZ, Varakunan S, Castellano D, Gutenkunst RN, Wahl LM. 2023. Shifts in mutation bias promote mutators by altering the distribution of fitness effects. The American Naturalist. 202:503–518.

Uecker H. 2017. Evolutionary rescue in randomly mating, selfing, and clonal populations. Evolution. 71:845–858.

Uecker H, Otto SP, Hermisson J. 2014. Evolutionary rescue in structured populations. The American Naturalist. 183:E17–E35.

Uphoff S, Kapanidis AN. 2014. Studying the organization of dna repair by single-cell and single-molecule imaging. DNA repair. 20:32–40.

van Velzen E. 2023. High importance of indirect evolutionary rescue in a small food web. Ecology Letters. 26:2110–2121.

Vanderwoude J, Azimi S, Read TD, Diggle SP. 2024. The role of hypermutation and collateral sensitivity in antimicrobial resistance diversity of pseudomonas aeruginosa populations in cystic fibrosis lung infection. Mbio. 15:e03109–23.

Vázquez-Mendoza I, Rodríguez-Torres EE, Ezadian M, Wahl LM, Gerrish PJ. 2024. Estimating the rate of mutation to a mutator phenotype. Axioms. 13:117.

Woodford N, Ellington MJ. 2007. The emergence of antibiotic resistance by mutation. Clinical Microbiology and Infection. 13:5–18.

Wu Y, Saddler CA, Valckenborgh F, Tanaka MM. 2014. Dynamics of evolutionary rescue in changing environments and the emergence of antibiotic resistance. Journal of theoretical biology. 340:222–231.

Wylie CS, Ghim CM, Kessler D, Levine H. 2009a. The fixation probability of rare mutators in finite asexual populations. Genetics. 181:1595–1612.

Wylie CS, Ghim CM, Kessler D, Levine H. 2009b. The fixation probability of rare mutators in finite asexual populations. Genetics. 181:1595–1612.

Xu K. 2023. Population rescue through an increase in the selfing rate under pollen limitation: Plasticity versus evolution. The American Naturalist. 202:337–350.

Xu K, Vision TJ, Servedio MR. 2023. Evolutionary rescue under demographic and environmental stochasticity. Journal of Evolutionary Biology. 36:1525–1538.

