## Supplementary text and figures for "The Role of Environmental Stress in Promoting Mutators Through Evolutionary Rescue: Quantitative Predictions"

### Online Supplement:
### Evolutionary Rescue Promotes Mutators,
### *GENETICS*

Marwa Tuffaha[1]
and
Lindi M. Wahl[1*]

1. Department of Mathematics, Western University, London, Ontario N6A 5B7, Canada;

### S1 Single Mutators

#### *S1.1 Analytical Methods*

##### *S1.1.1 Probability of Rescue by the Wildtype*

Let $p_i(t)$ be the probability that $i$ rescue mutants with wildtype background are present at time $t$ (sometimes written as $p_i$ in the equations to follow). The pgf describing the number of rescue mutants at time $t$ is

$$\phi_W(x,t) = \sum_{i=0}^{\infty} p_i(t)x^i. \tag{S1}$$

Selection pressure due to competition affects the rescue mutants through their division rates $f_{R_W}(t)[1-N(t)/K]$, where $N(t)$ is the total population size. Assuming this rescuing subpopulation is sufficiently rare, $N(t)$ can be approximated by the sum of the wildtype population, $N_W(t)$, and the expected number of mutator individuals, $\mathbb{E}[N_M]$, so that the rescue mutant's effective growth rate is $f_{R_W}^{\text{eff}}(t) = f_{R_W}(t)[1-(N_W(t)+\mathbb{E}[N_M(t)])/K]$.

Within a $\Delta t$ increment in time, the change in the pgf $\phi_W$ comes from two potential events that can happen in the time interval $[t, t+\Delta t]$; a rescue mutant either replicates or dies. Assuming the rescue mutant originating from the wildtype has the same death rate as the wildtype, $g$, the equation describing these events is given as follows:

$$\phi_W(x,t+\Delta t) - \phi_W(x,t) = \sum ip_i f_{R_W}^{\text{eff}}(t)\Delta t x^{i+1} - \sum ip_i f_{R_W}^{\text{eff}}(t)\Delta t x^i + \sum ip_i g\Delta t x^{i-1} - \sum ip_i g \Delta t x^i. \tag{S2}$$

Simplifying, we find that the time evolution of $\phi_W$ is then given by

$$\frac{\partial \phi_W}{\partial t} = (1-x)(g - f_{R_W}^{\text{eff}}(t)x)\frac{\partial \phi_W}{\partial x}. \tag{S3}$$

This PDE is equivalent to the one presented by Marrec and Bitbol (2020) (except for the definition of the effective growth rate), where assuming one mutant emerges at time $t_0$, as the authors also found, the method of characteristics can be used to find an explicit solution given by

$$\phi_W(x,t) = 1 + \left( \frac{e^{\rho(t)}}{x-1} - \int_{t_0}^{t} f_{R_W}^{\text{eff}}(u)e^{\rho(u)}du \right)^{-1}, \tag{S4}$$

$$\text{where} \qquad \rho(t) = \int_{t_0}^{t} (g - f_{R_W}^{\text{eff}}(u))du. \tag{S5}$$

As with any pgf (Allen, 2015), the extinction probability is then the value of the pgf evaluated at $x = 0$ as $t \to \infty$, and the fixation probability is the complement of that, $P_{\text{fix}}^W(t_0) = 1 - \lim_{t\to\infty} \phi_W(0,t)$. As found by Marrec and Bitbol (2020), the fixation probability is:

$$P_{\text{fix}}^W(t_0) = \frac{1}{1 + g\int_{t_0}^{\infty} e^{\rho(t)}dt}, \tag{S6}$$

where we can use

$$\int_{t_0}^{t} (g - f_{R_W}^{\text{eff}}(u))e^{\rho(u)}du = e^{\rho(t)} - 1. \tag{S7}$$

Plugging eq. (S6) into eq. (9) in the main text gives the probability of rescue by the wildtype.

##### *S1.1.2 Probability of Rescue by the Mutator*

Let $p_{ij}(t)$ be the probability that $i$ mutator individuals and $j$ rescue individuals with mutator background are present at time $t$ (sometimes written as $p_{i,j}$ in the equations to follow). The multitype pgf describing the mutator lineage is

$$\phi_M(x,y,t) = \sum_{i=0}^{\infty}\sum_{j=0}^{\infty} p_{ij}(t)x^i y^j, \tag{S8}$$

where the dummy variables $x$ and $y$ represent the mutator and its rescue mutant, respectively.

Again, assuming the rescuing populations are rare, we can approximate the population size by $N_W(t) + \mathbb{E}[N_M(t)]$ and use effective growth rate functions for the mutator and its rescue mutant, $f_M^{\text{eff}}(t) = f_M(t)[1-(N_W(t)+\mathbb{E}[N_M(t)])/K]$ and $f_{R_M}^{\text{eff}}(t) = f_{R_M}(t)[1-(N_W(t)+\mathbb{E}[N_M(t)])/K]$, respectively.

Within a $\Delta t$ increment in time, the change in this multitype pgf comes from five potential events that can happen in the time interval $[t, t+\Delta t]$. A mutator replicates and mutates at the rescue allele, replicates without a mutation, or dies, while a rescue mutant can only either replicate or die. Assuming the mutator's rescue mutants also have death rate, $g$, the equation describing these events is given as follows, where each row corresponds to one of the mentioned events, respectively:

$$\begin{aligned}\phi_M(x,y,t+\Delta t) - \phi_M(x,y,t) = &\sum\sum ip_{ij}f_M^{\text{eff}}(t)\mu'\Delta t x^i y^{(j+1)} - \sum\sum ip_{ij}f_M^{\text{eff}}(t)\mu'\Delta t x^i y^j \\ &+ \sum\sum ip_{ij}f_M^{\text{eff}}(t)(1-\mu')\Delta t x^{i+1}y^j - \sum\sum ip_{ij}f_M^{\text{eff}}(t)(1-\mu')\Delta t x^i y^j \\ &+ \sum\sum ip_{ij}g\Delta t x^{i-1}y^j - \sum\sum ip_{ij}g\Delta t x^i y^j \\ &+ \sum\sum jp_{ij}f_{R_M}^{\text{eff}}(t)\Delta t x^i y^{j+1} - \sum\sum jp_{ij}f_{R_M}^{\text{eff}}(t)\Delta t x^i y^j \\ &+ \sum\sum jp_{ij}g\Delta t x^i y^{j-1} - \sum\sum jp_{ij}g\Delta t x^i y^j \ .\end{aligned} \tag{S9}$$

The time evolution of the pgf is then given by

$$\frac{\partial \phi_M}{\partial t} = \Big[(1-x)g + f_M^{\text{eff}}(t)x[(1-\mu')x + \mu' y - 1]\Big]\frac{\partial \phi_M}{\partial x} + (1-y)(g - f_{R_M}^{\text{eff}}(t)y)\frac{\partial \phi_M}{\partial y}. \tag{S10}$$

This PDE describes the fate of an initially rare mutator population. Initially, we assume that only one mutator individual emerges at time $t_0$ and no rescue mutant, i.e.,

$$\phi_M(x,y,t_0) = x. \tag{S11}$$

To integrate eq. (S10) on a time interval $t \in [t_0, T]$, we use the method of characteristics to write it as a system of ODEs:

$$\text{Mutator: } \frac{dx}{d\tau} = (1-x)g + f_M^{\text{eff}}(T-\tau)x[(1-\mu')x + \mu' y - 1], \tag{S12}$$

$$\text{Rescue Mutants: } \frac{dy}{d\tau} = (1-y)(g - f_{R_M}^{\text{eff}}(T-\tau)y), \tag{S13}$$

$$\text{Time to mutation: } \frac{dt}{d\tau} = -1. \tag{S14}$$

The time variable in the system (S12-S14), $\tau$, is defined in terms of the time variable in the PDE (S10), $t$, as $\tau = T - t$, so that the time interval for the ODE system is $\tau \in [0, T - t_0]$. An explicit solution to eq. (S13) is possible, but the use of that solution does not allow for an explicit solution to eq. (S12), and thus we integrate this system numerically (ode45, Matlab©).

Recall that in the method of characteristics, to evaluate the pgf $\phi_M$ at any particular $x$, $y$ and $T$ values, we use $x$, $y$ and $T$ as initial conditions and integrate system (S12-S14) backwards in time until $t = t_0$. The value of $\phi_M(x, y, T)$ is then equal to $\phi_M$ evaluated at the resulting coordinates, that is, $\phi_M(x(T - t_0), y(T - t_0), t_0)$. Since the value of $\phi_M$ at $t = t_0$ is known (eq. (S11)), the desired value $\phi_M(x, y, T)$ is simply given by $x(T - t_0)$.

Our goal is to estimate the extinction probability of the mutator lineage, i.e., to evaluate $\phi_M(0, 0, T)$ in the limit as $T$ approaches infinity. This corresponds to taking $x(\tau = 0) = y(\tau = 0) = 0$ as initial conditions in system (S12-S14). We increase the value of $T$ until the solution converges, which gives us the extinction probability, $P^M_{\text{ext}}(t_0)$, as the value of the $x$ variable as per eq. (S11). Then, the probability of fixation of the lineage of this mutator is given by $P^M_{\text{fix}}(t_0) = 1 - P^M_{\text{ext}}(t_0)$. Using $P^M_{\text{fix}}(t_0)$ in eq. (**??**) gives the probability of rescue by the mutator.

#### *S1.2 Simulation Methods*

We use a Gillespie algorithm (Gillespie, 1977), extending the code generated by Marrec and Bitbol (Marrec and Bitbol, 2020) and using the same assumption that the time interval between any two consecutive events is small and thus the growth rate changes in that time interval can be ignored. So, the values of the growth rate functions to be evaluated at any event are taken at the time $t$ of the last event that occurred.

We consider four populations: the wildtype ($W$), the mutator ($M$), the rescue mutants founded by the wildtype ($R_W$), and rescue mutants founded by the mutator ($R_M$). Upon division, the probability of mutation from W to M is $\mu_M$, from W to $R_W$ is $\mu$, and from M to $R_M$ is $\mu' = F\mu$. The populations are subject to a carrying capacity $K$ and the total population size at time $t$, $N(t)$, is the sum of all these four populations. We have 11 elementary events that can occur which are:

- $W \to 2W$: Replication without mutation of a wildtype individual with rate $h^+_W = f_W(t)(1 - N(t)/K)(1 - \mu_M - \mu)$.
- $W \to W + M$: Replication with mutation of a wildtype individual to a mutator with rate $h_{WM} = f_W(t)(1 - N(t)/K)\mu_M$.
- $W \to W + R_W$: Replication with mutation of a wildtype individual at the rescue allele with rate $h_{WR} = f_W(t)(1 - N(t)/K)\mu$.
- $W \to \emptyset$: Death of a wildtype individual with rate $h^-_W = g_W$.
- $M \to 2M$: Replication without mutation of a mutator individual with rate $h^+_M = f_M(t)(1 - N(t)/K)(1 - \mu')$.
- $M \to M + R_M$: Replication with mutation of a mutator individual at the rescue allele with rate $h_{MR} = f_M(t)(1 - N(t)/K)\mu'$.
- $M \to \emptyset$: Death of a mutator individual with rate $h^-_M = g_M$.

- $R_W \to 2R_W$: Replication of a rescue mutant founded by the wildtype with rate $h^+_{R_W} = f_{R_W}(t)(1 - N(t)/K)$.
- $R_W \to \emptyset$: Death of a rescue mutant founded by the wildtype with rate $h^-_{R_W} = g_{R_W}$.
- $R_M \to 2R_M$: Replication of a rescue mutant founded by the mutator with rate $h^+_{R_M} = f_{R_M}(t)(1 - N(t)/K)$.
- $R_M \to \emptyset$: Death of a rescue mutant founded by the mutator with rate $h^-_{R_M} = g_{R_M}$.

The total rate of events is

$$H = (h^+_W + h_{WM} + h_{WR} + h^-_W)N_W + (h^+_M + h_{MR} + h^-_M)N_M + (h^+_{R_W} + h^-_{R_W})N_{R_W} + (h^+_{R_M} + h^-_{R_M})N_{R_M}. \quad \text{(S15)}$$

Simulation steps are as follows:

1. Initialization: For mutators coming from *de novo* mutation, the populations start at zero initial values except for the wildtype, which starts at $N^0_W = (1 - g_W/f_W(0))K$. On the other hand, for pre-existing mutators, we set $N^0_W = (1 - \alpha)(1 - g_W/f_W(0))K$ and $N^0_M = \alpha(1 - g_W/f_W(0))K$, while the rescue mutant populations start at zero. If $f_W(0) = 0$, which can happen at mutation rates higher than or equal to the critical value leading to mutational meltdown, $\nu_c$, then no populations exist and the rescue probability is of course zero.
2. We randomly sample the time increment $dt$ from an exponential distribution with mean $1/H$. With probabilities $h/H$ of the event, proportional to each event's rate $h$, we randomly choose the next event to occur.
3. We increase the time to $t = t + dt$ and update the growth rate function values and the populations according to the event chosen at step 2.
4. We iterate steps 2 and 3 until the wildtype population goes extinct ($N_W = 0$), where we count this simulation trial as a failure, or one of the rescue mutant populations fixes and reaches the carrying capacity ($N_{R_W} > K$ or $N_{R_M} > K$), where we count this simulation trial as a success.

The simulation is repeated $q$ times and the probability of rescue is given by the number of successes divided by $q$. In fact, the simulation described above can be used to find four different probabilities:

- The probability of rescue by the wildtype in the absence of mutators, $Pr_{W_0}$ (eq. 7 when $\mu_M$ is set to zero), can be found by setting $\mu_M = 0$ to turn off the mutator population.
- The probability of rescue by the wildtype in the presence of mutators, $Pr_W$ (eq. 7), can be found by setting $\mu' = 0$ to turn off the rescue by the mutator.
- The probability of rescue by the mutator only, $Pr_M$ (eq. 15), can be found by setting $\mu = 0$ to turn off the rescue by the wildtype.
- The probability of rescue by either the wildtype or the mutator, $Pr_{all}$ (eq. 23), can be found by turning on all the three mutations, wildtype to mutator, wildtype to rescue, and mutator to rescue.

#### *S1.3 Supplementary Results*

##### *S1.3.1 Time Series*

To provide greater insight into the temporal dynamics of evolutionary rescue, we present time series plots of the wildtype and mutator populations across different parameter scenarios (Figures S1-S5). The plotted WT populations in blue represent the sums original WT ($W$) and its rescue mutants ($R_W$), and similarly, the plotted mutator populations in red represent the sum of ($M$) and ($R_M$). These plots illustrate how the emergence and dynamics of $R_M$ and $R_W$ lineages vary under different rates of environmental change (Hill coefficient $n = 2$ vs. $n = 10$) and under conditions of *de novo* emergence vs. pre-existing mutators.

The trajectories of 5000 replicates are depicted in figures S1-S5 as semi-transparent lines to illustrate variability across simulations, and the means are shown with solid thick lines. In the *de novo* emergence cases, the mutator emergence factor $E = 500$. Pre-existing mutators in these figures start at a fraction $\alpha = 5\%$ of the population. Other parameters are $F = 5$, $\nu = 0.01$, $g = 0.1$ and $\theta = 500$. In the absence of environmental change, simulations have only one outcome (fig. S1). When environmental stress in introduced, the population either goes extinct, is rescued by the WT, or is rescued by the mutator. In figures S2-S5, we categorize the instances based on each of the three outcomes and plot them separately for clarity and ease of interpretation.

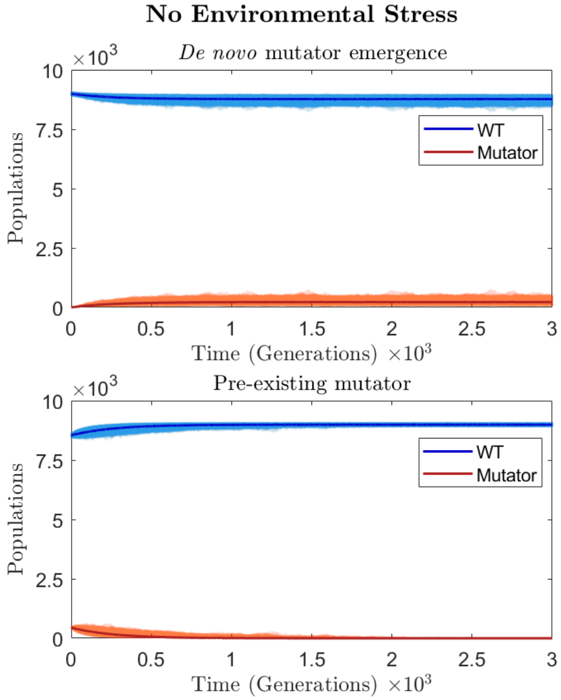


Figure S1: Trajectories of the WT and mutator populations in the absence of environmental stress. Co-existence occurs in the case of *de novo* mutator emergence (upper panel), while mutators go extinct if they only pre-existed (lower panel). Parameters are explained in section S1.3.1.

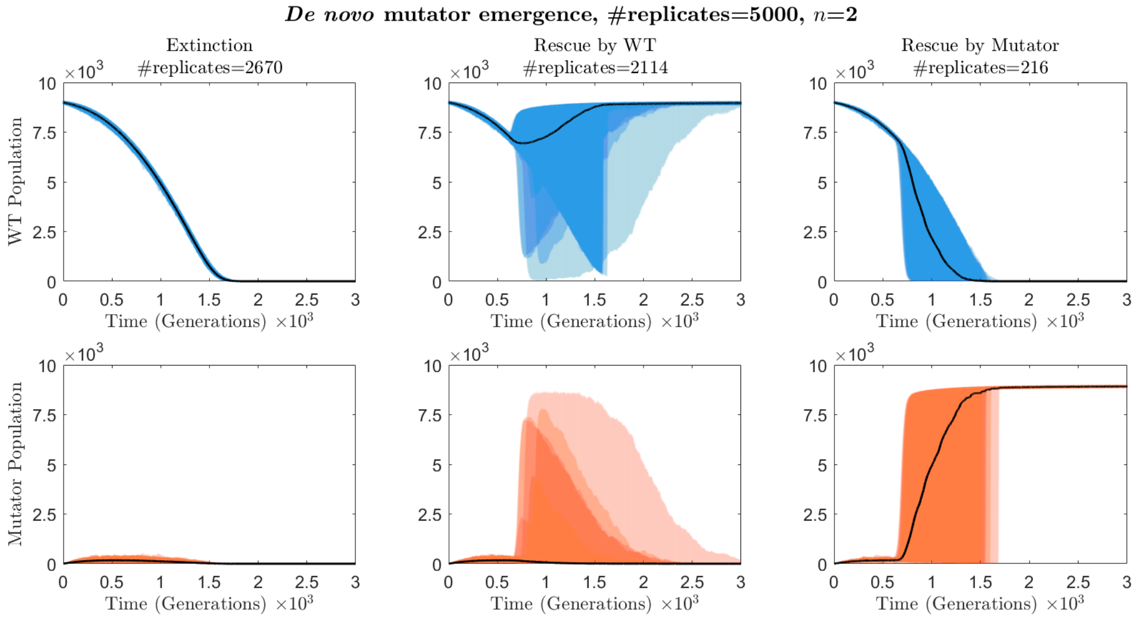


Figure S2: Trajectories of the WT and *de novo* mutator populations in the case of slow environmental change ($n = 2$), separated by outcome. Parameters are explained in section S1.3.1.

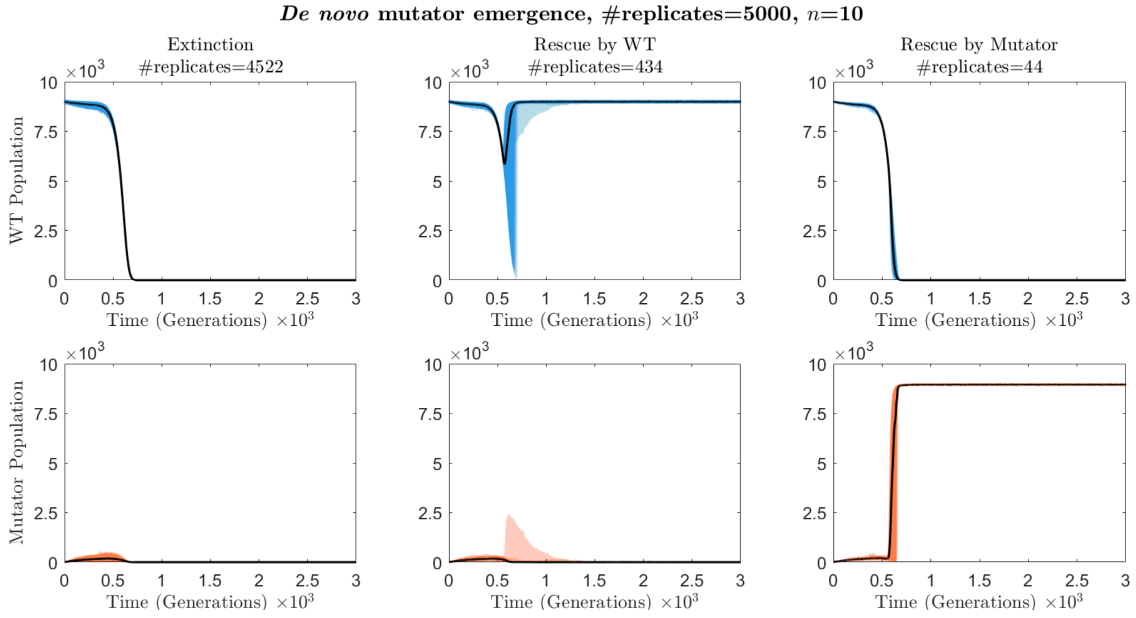


Figure S3: Trajectories of the WT and *de novo* mutator populations in the case of fast environmental change ($n = 10$), separated by outcome. Parameters are explained in section S1.3.1.

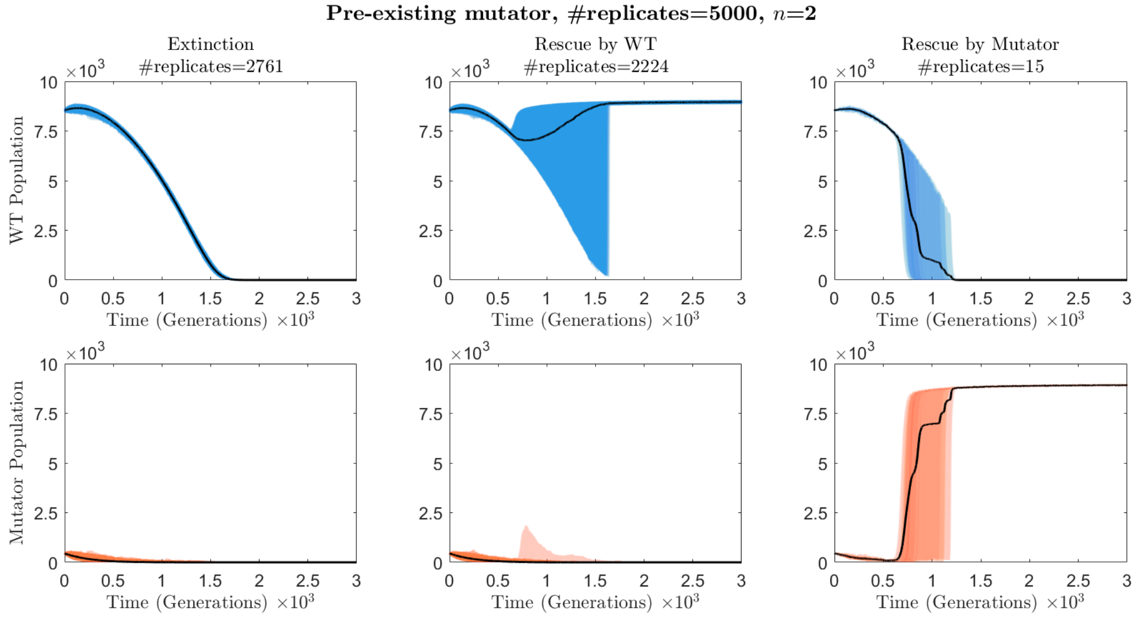


Figure S4: Trajectories of the WT and pre-existing mutator populations in the case of slow environmental change ($n = 2$), separated by outcome. Parameters are explained in section S1.3.1.

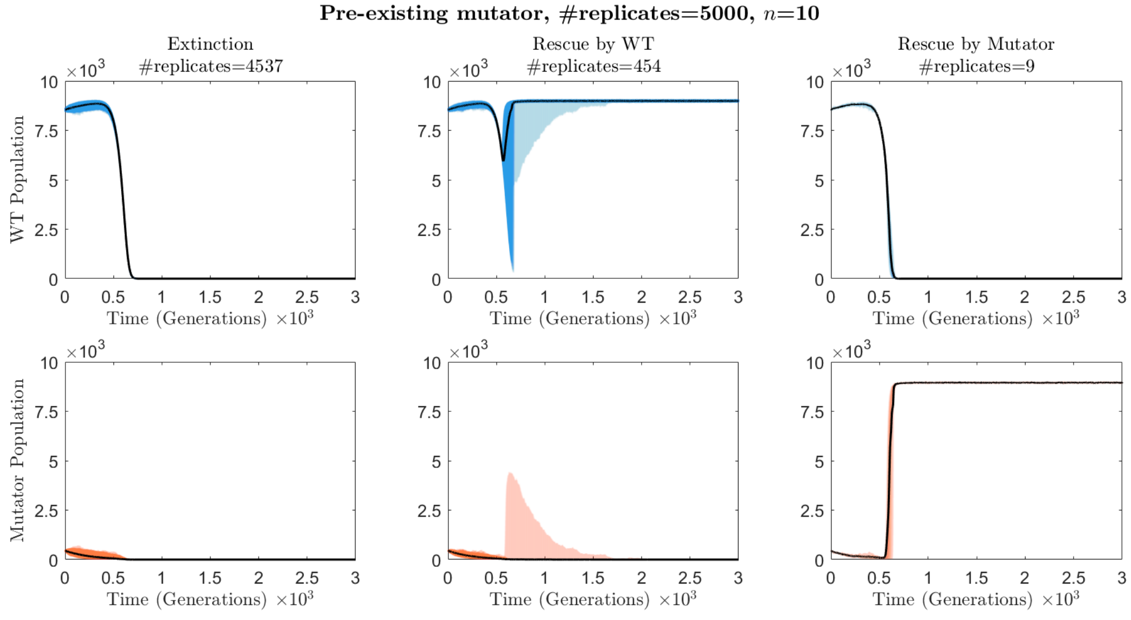


Figure S5: Trajectories of the WT and pre-existing mutator populations in the case of fast environmental change ($n = 10$), separated by outcome. Parameters are explained in section S1.3.1.

##### *S1.3.2 The Centered Time of Environmental Change has Different Effects whether Mutators Emerge or Pre-exist*

Early environmental changes (low $\theta$) give low chances for mutator *de novo* emergence, decreasing in turn the probability of rescue by mutators (fig. S6A,B). In contrast, when mutators pre-exist, they decline through time and thus earlier environmental changes are expected to be advantageous to the mutator, giving it more chance to rescue the population. Nonetheless, we see a slight increase in the highest possible probability reached as $\theta$ decreases from 500 to 50, with a peak shift towards favoring higher wildtype mutation rates (fig. S6C,D). Decreasing $\theta$ more keeps shifting the peak at the same direction, but its height gets lower, indicating that when the number of generations prior to $\theta$ becomes too low, the rescuing mutants get fewer chances of occurring before the environmental change eliminates the mutator population (fig. S6C,D).

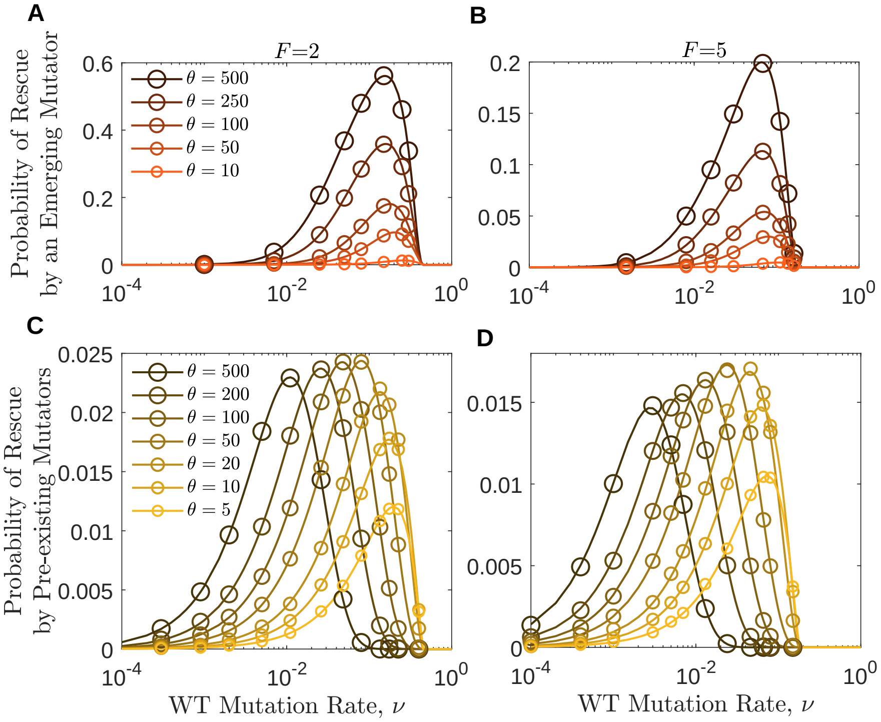


Figure S6: The time of environmental change has different effects whether mutators emerge or pre-exist. A,B: Emerging mutators with $E = 500$ have lower probabilities of rescue for earlier environmental changes. C,D: Pre-existing mutators at a $\alpha = 5\%$ frequency are more likely to rescue the population for higher wildtype mutation rates when the central time of the environmental change is early, but more likely to rescue a population founded by a wildtype with intermediate mutation rates for later environmental changes. Mutator strength is $F = 2$ in the left panels, and $F = 5$ in the right panels. Other parameters are $n = 2$ and death rates are 0.1. Circles show simulation results from over $25 \times 10^4$ replicates for emerging mutators and over $45 \times 10^4$ replicates for pre-existing mutators.

For completeness, in fig. S7 we also show analytical results for large values of $\theta$. For pre-existing mutators, the population experiences a gradual decline in mutator frequency over time due to their selective disadvantage, as no additional mutators are introduced and selection acts to purge them (fig. S1). In the absence of ongoing mutational input, this results in an eventual loss of mutators from the population, except for a small fraction that may transiently persist for a long time due to genetic drift. Consequently, the rescue probabilities converge to zero more slowly once the environmental change occurs after a sufficiently long delay (lower panels). For *de novo* mutations, because our model does not include back mutations, lineages carrying the rescue mutation gradually accumulate during these extended time intervals, leading to an overestimation of the rescue probability (upper panels). This suggests that incorporating back mutation would be important in future models aiming to capture the dynamics accurately over long evolutionary timescales.

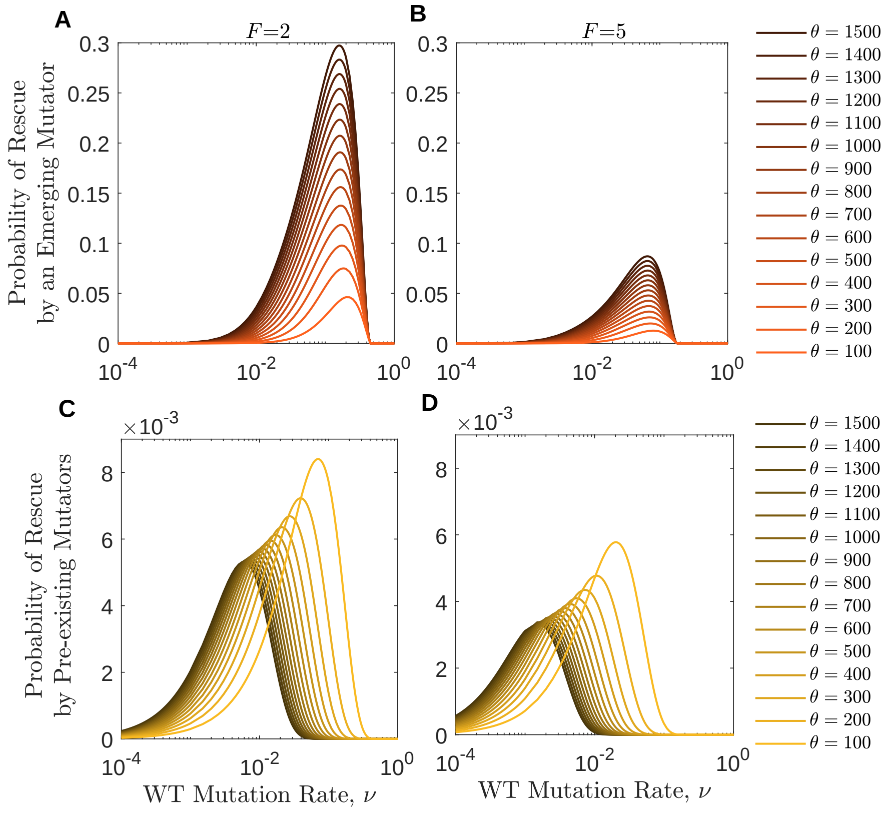


Figure S7: Rescue probabilities stabilize with large $\theta$ in the pre-existing case, but are overestimated in the *de novo* case without back mutation. A,B: Emerging mutators with $E = 500$ have higher probabilities of rescue for later environmental changes due to the accumulation of rescue mutants on mutator background. C,D: Probabilities for pre-existing mutators at $\alpha = 5\%$ initial frequency decrease slowly due to purging by selection in the absence of ongoing input, with only rare persistence via drift. Mutator strength is $F = 2$ in the left panels, and $F = 5$ in the right panels. Other parameters are $n = 10$ and $g = 0.1$.

##### *S1.3.3 Post-Rescue Mutator Switching (Allowing $R_W \to R_M$ mutations)*

While the main simulations and analytical work did not include the possibility of a wildtype rescue mutant (RW) subsequently acquiring a mutator allele, we ran a small number of supplementary simulations to assess the potential impact of this pathway. Figure S8 compares simulation outcomes with and without $R_W \to R_M$ mutations under a representative parameter set.

We observed no difference in the probability of rescue by the mutator ($R_M$) between the two scenarios. This suggests that allowing mutator acquisition after a rescue mutation does not materially alter the evolutionary dynamics, likely due to the low frequency of $R_W$ and the limited benefit conferred by switching to a mutator state post-rescue.

As such, in addition to the analytical complication in including these mutations, we chose to retain our original simulation and analytical design without $R_W \to R_M$ mutations, but include this supplementary result for completeness.

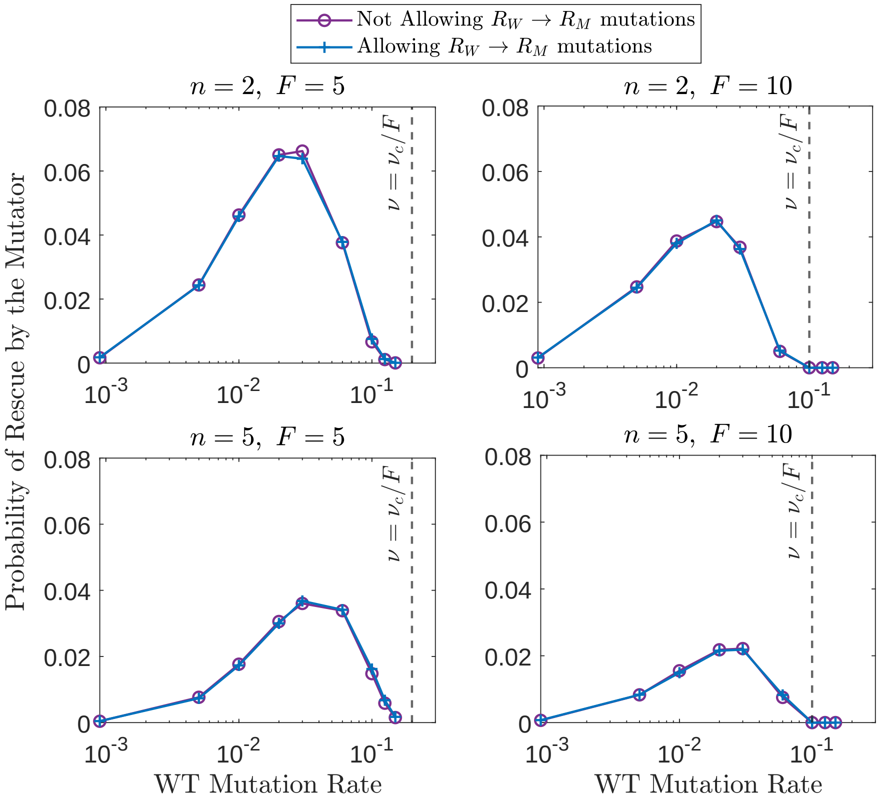


Figure S8: Comparison of the probability of rescue by mutators ($R_M$) with and without allowing post-rescue mutator acquisition ($R_W \to R_M$ mutations). Across the tested parameter values, no difference was observed between the two conditions.

#### *S1.4 Fraction Rescued by Mutator*

In addition to mutator advantage (eq. 24), another useful measure of the advantage of mutator evolution or pre-existence in the evolutionary rescue process is the fraction of rescue events attributed to mutators out of all instances in which the population is rescued. Deriving an analytical estimate for this fraction presents significant challenges due to the dynamic nature of the process.

One major complication is that WT-driven and mutator-driven rescue events do not occur uniformly throughout time. Since mutators arise from WT mutations, their rescue events tend to take place later in the evolutionary timeline. This non-uniform distribution means that standard analytical approximations, which typically assume constant rates, would likely overestimate the fraction of rescues attributed to mutators. Furthermore, in scenarios where mutators are pre-existing, our system is not at equilibrium when environmental change begins. The timing of the environmental shift greatly influences whether rescue occurs through WT or mutator lineages, further complicating an analytical approach.

Due to these complexities, we only show simulation results for the fraction of mutator-driven rescue events (fig. 3). Simulations allow for an accurate representation of time-dependent mutation and selection processes without oversimplifying critical temporal effects. While future studies may develop analytical approximations of the time distributions of these rescue events, allowing to account for these challenges, such an approach is beyond the scope of this study.

##### *S1.5 Alternative Mutator Model: Additive Increase in Mutation Rate*

In the main analysis, we define the mutation rate of mutator lineages as a constant multiple of the wildtype mutation rate, $\nu' = F\nu$, following standard theoretical and empirical practice. However, as discussed in the main text, alternative relationships may arise in biological systems, such as additive increases under a modular repair architecture. To illustrate the flexibility of our framework, we implemented an additive formulation where $\nu' = \nu + \Delta$ and compared the resulting outcomes to those from the multiplicative model. The results (Figure S9) show that while the general trends remain similar, the mutator's chance of rescuing the population increases with $\Delta$ at low WT mutation rates, but is almost similar at high WT mutation rates (Figure S9C). The flipping effect (as in fig. 5 in the main text) is still there but is barely seen as the probability keeps increasing for all chosen values of $\Delta$ even at high WT mutation rates since the cost is not amplified in this region anymore. Another observation is that the mutator advantage saturates at low mutation rates as $\nu$ becomes negligible compared to $\nu'$ (Figure S9B). This comparison highlights the importance of carefully considering how mutator effects are modeled when interpreting their evolutionary consequences.

#### S2 Multiple Mutators Emergence

##### *S2.1 Analytical Predictions*

In the main text, we assume a single mutator population with an $F$-fold increase in mutation rate that emerges at rate $\mu_M$. In reality, a declining population might have multiple mutators that can emerge and rescue it.

Let $m$ be the number of potential mutators, each with an $F_i$-fold higher mutation rate than the wildtype, i.e., the $i$-th mutator mutates at rate $\nu'_i = F_i\nu$ per genome per generation but only at the rescue allele at rate $\mu'_i = F_i\mu$, where $i = 1, .., m$. If the $i$-th mutator has an emergence rate $\mu_{M_i}$, the overall emergence rate of mutators is $\mu_M = \sum_{i=1}^{m} \mu_{M_i}$. In this case, even when $\mu_M$ is large, if the emergence rates of the individual mutators, $\mu_{M_i}$, are small enough, the populations of

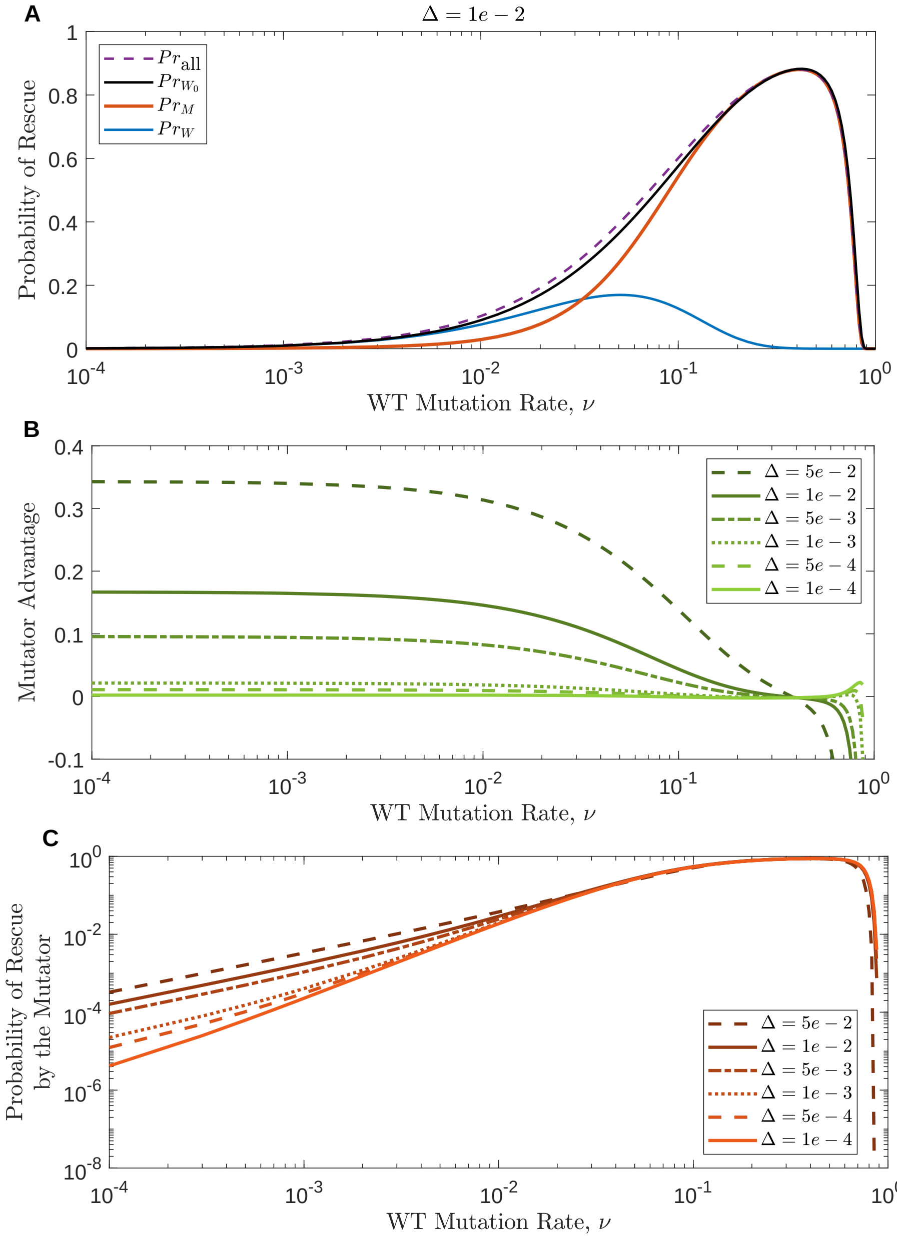


Figure S9: Results of the additive model, where the mutator mutation rate $\nu' = \nu + \Delta$. Parameter values are $n = 10$, $\theta = 500$, $E = 2000$ and $g = 0.1$.

the mutators will be rare and the above approximation can be used to estimate the probability of rescue by one, some, or all of these mutators.

The wildtype population will still be governed by eq. (5), where the competition between the wildtype and the mutators is considered by allowing the expected value of the total mutator population, $\mathbb{E}[N_M]$, to compete with the wildtype. In this case, $\mathbb{E}[N_M] = \sum_{i=1}^{m} \mathbb{E}[N_{M_i}]$ is the sum of the expected population sizes of the mutator subpopulations, which are governed by the following $m$ ODEs (similar to eq. 6):

$$\frac{d\mathbb{E}[N_{M_i}]}{dt} = \mu_{M_i} f_W(t) \left(1 - \frac{N_W + \mathbb{E}[N_M]}{K}\right) N_W + \left[f_{M_i}(t)\left(1 - \frac{N_W + \mathbb{E}[N_M]}{K}\right) - g_{M_i}\right] \mathbb{E}[N_{M_i}], \tag{S16}$$

where the $i$-th mutator has a death rate $g_{M_i}$, and its growth rate function $f_{M_i}(t)$ is given by replacing $\nu'$ with $\nu'_i$ in eq. (3).

The branching process now needs to be applied for each mutator alone to get its own probability of rescue, where each mutator now independently emerges from the wildtype population. For the $i$-th mutator, the PDE to solve will be the same as eq. (18) but replacing the parameters and the growth rate function to match the parameters of that mutator as follows:

$$\frac{\partial \phi_{M_i}}{\partial t} = \left[(1-x)g_{M_i} + f^{\text{eff}}_{M_i}(t)x[(1-\mu'_i)x + \mu'_i y - 1]\right] \frac{\partial \phi_{M_i}}{\partial x} + (1-y)(g_{R_{M_i}} - f^{\text{eff}}_{R_{M_i}}(t)y)\frac{\partial \phi_{M_i}}{\partial y}. \tag{S17}$$

The numerical solution of the corresponding system of the method of characteristics gives the extinction probability of that mutator, whose complement will be its fixation probability $P^{M_i}_{\text{fix}}(t_0)$. The probability of rescue by the $i$-th mutator is then given by

$$Pr_{M_i} = \mu_{M_i} \int_0^\infty P^{M_i}_{\text{fix}}(u) N_W(u) f_W(u) \left(1 - \frac{N_W(u) + \mathbb{E}[N_M(u)]}{K}\right) du. \tag{S18}$$

If the population is rescued by any of the $m$ mutators, it cannot be rescued by any other one, and thus the probability of rescue by any mutator is given by:

$$Pr_M = 1 - \prod_{i=1}^{m}(1 - Pr_{M_i}). \tag{S19}$$

This approach assumes no competition between the multiple mutator populations, which is appropriate when each mutator population is small, but as they grow large, this approximation becomes less accurate. This assumption is relaxed in the simulations.

#### *S2.2 Simulation Methods*

If we want $m$ mutator populations to emerge from the wildtype, each with an emergence rate $\mu_{M_i}$, an elevated mutation rate at the rescue allele $\mu'_i$, and death rate $g_{M_i}$, then we will have more subpopulations and more events happening in the simulation. In fact, there will be $m$ mutator populations, $N_{M_i}$, and $m$ rescue population each founded by one of the mutators, $N_{R_{M_i}}$, for $i = 1, .., m$. The total population size is now $N(t) = N_W(t) + N_{R_W}(t) + \sum_{i=1}^{m}(N_{M_i} + N_{R_{M_i}})$.

Letting $\mu_M = \sum_{i=1}^{m} \mu_{M_i}$ be the overall emergence rate of mutators, five events will stay the same: these are the wildtype replication without mutation $W \to 2W$, the wildtype replication with mutation at the rescue allele $W \to W + R_W$, the wildtype death $W \to \emptyset$, the replication of a rescue mutant founded by the wildtype $R_W \to 2R_W$, and the death of a rescue mutant founded by the wildtype $R_W \to \emptyset$. However, each of the other six events will now be replaced my $m$ similar events for the different mutators as follows:

- $W \to W + M_i$: Replication with mutation of a wildtype individual to the $i$-th mutator with rate $h_{WM_i} = f_W(t)(1 - N(t)/K)\mu_{M_i}$.
- $M_i \to 2M_i$: Replication without mutation of an individual from the $i$-th mutator population with rate $h^+_{M_i} = f_{M_i}(t)(1 - N(t)/K)(1 - \mu'_i)$.
- $M_i \to M_i + R_{M_i}$: Replication with mutation at the rescue allele of an individual from the $i$-th mutator population with rate $h_{M_iR} = f_{M_i}(t)(1 - N(t)/K)\mu'_i$.
- $M_i \to \emptyset$: Death of an individual from the $i$-th mutator population with rate $h^-_{M_i} = g_{M_i}$.
- $R_{M_i} \to 2R_{M_i}$: Replication of a rescue mutant founded by the $i$-th mutator with rate $h^+_{R_{M_i}} = f_{R_{M_i}}(t)(1 - N(t)/K)$.
- $R_{M_i} \to \emptyset$: Death of a rescue mutant founded by the $i$-th mutator with rate $h^-_{R_{M_i}} = g_{R_{M_i}}$.

The total rate of events is

$$\begin{aligned}\bar{H} =&(h^+_W + \sum_{i=1}^{m}(h_{WM_i}) + h_{WR} + h^-_W)N_W + \sum_{i=1}^{m}[(h^+_{M_i} + h_{M_iR} + h^-_{M_i})N_{M_i}] \\ &+(h^+_{R_W} + h^-_{R_W})N_{R_W} + \sum_{i=1}^{m}[(h^+_{R_{M_i}} + h^-_{R_{M_i}})N_{R_{M_i}}]. \end{aligned} \tag{S20}$$

We follow the same steps described in the previous simulation considering all the $6m+5$ events and counting the trial as a success if any of the rescue mutants, whether from the wildtype or a mutator background, reaches the carrying capacity.

It is also possible to decide which probability to find by

- either turning off all mutator emergence rates $\mu_{M_i} = 0$ to get the probability of rescue by the wildtype in the absence of mutators, $Pr_{W0}$ (eq. 7 when $\mu_M$ is set to zero),
- or turning off rescue by the mutators $\mu'_i = 0$ to get the probability of rescue by the wildtype in the presence of mutators, $Pr_W$ (eq. 7),
- or turning off rescue by the wildtype only $\mu = 0$ to get the probability of rescue by the overall mutator population, $Pr_M$ (eq. 15).
- or turning on all mutations to get the probability of rescue by either the wildtype or a mutator, $Pr_{all}$ (eq. 23),

#### *S2.3 Parameter Values*

We arbitrarily choose to have $m = 20$ mutator subpopulations with an overall mutators emergence factor $e = 2,000$ so that the total mutators emergence rate $\mu_M = 2,000\mu$, which is (for simplicity) divided equally among the mutators so that the $i$-th mutator has an emergence rate $\mu_{M_i} = \mu_M/m$. Also, the mutation rate multiple $F_i$ for the $i$-th mutator is chosen from the set

$$F_{\text{vals}} = \{2, 5, 10, 15, 20, 25, 30, 35, 40, 45, 50, 55, 60, 65, 70, 75, 80, 85, 90, 100\}.$$

We allow repetition in choosing the mutation rate multiples from $F_{\text{vals}}$ in the 20 mutator subpopulations; for example, two mutator subpopulations $i$ and $j$ can have the same mutation rates $\nu'_i = \nu'_j = \bar{F}\nu$ if $F_i = F_j = \bar{F}$, which corresponds to one mutator subpopulation with a mutation rate multiple $\bar{F}$ and an emergence rate $\mu_{M_i} + \mu_{M_j} = 2\mu_M/m$. If the same $\bar{F}$ value is repeated $\bar{m}$ times, this corresponds to one mutator subpopulation with emergence rate $\bar{\mu}_M = \bar{m}\mu_M/m$.

We consider different combinations for the $F_i$ values. As a baseline case, we let each mutator subpopulation have a different mutation rate, meaning that there is no bias towards the emergence of any mutation rate more than any other. We refer to this case with the label $k = 0$, which we call the "bias towards larger mutation rate increases". From its name, positive values of $k$ mean that higher $F$ values are more likely to be chosen from the set $F_{\text{vals}}$, and vice versa for negative values of $k$. Table S1 shows the different combinations we choose and the $k$ value referring to each one of them. We emphasize that $k$ is not a computed parameter, but is simply a label to conveniently refer to these chosen test cases.

#### *S2.4 Results*

In the high wildtype mutation rate regime (right-hand side of figures), the weakest mutator subpopulation dominates the behavior of the probability of rescue by the overall population of mutators, and vice versa (fig. S10B). For example, in the scenarios we suggested (Table S1), since the lowest mutator strength we allow is $F = 2$, when a mutator with this strength is present in the mutators population ($k \leq 0$) it is the only mutator that can still have births and thus rescue the population up to high wildtype mutation rates that are close $\nu = \nu_c/2$, and thus the probability of rescue by mutators stays positive in these scenarios until that point, and this probability increases for higher fractions of this mutator's population (lower $k$). In the other scenarios where this mutator strength is not included ($k > 0$), the probability of rescue becomes zero when the wildtype mutation rate reaches the critical value for the weakest mutator present. In contrast, at low wildtype mutation rates, the higher the value of the strongest mutator present, and the larger its proportion in the population, the higher the probability of rescue by mutators.

We note that our analytical predictions in the multiple mutators case overestimate the rescue probability at high wildtype mutation rates, $\nu$, and when favoring mild mutator strengths, $k < 0$, since both factors lead to a lower overall mutation rate cost. This allows the mutator subpopulations to grow and compete with one another, while we do not consider this kind of competition in our analytical predictions.

### References

Allen, L. J. 2015. Stochastic population and epidemic models. Mathematical biosciences lecture series, stochastics in biological systems 1:120–128.

| $F_{\text{vals}}$ | $k = -3$ | $k = -2$ | $k = -1$ | $k = 0$ | $k = 1$ | $k = 2$ | $k = 3$ |
|---|---|---|---|---|---|---|---|
| 2 | 4 | 3 | 2 | 1 | 0 | 0 | 0 |
| 5 | 4 | 3 | 2 | 1 | 0 | 0 | 0 |
| 10 | 3 | 2 | 2 | 1 | 0 | 0 | 0 |
| 15 | 3 | 2 | 2 | 1 | 0 | 0 | 0 |
| 20 | 2 | 2 | 2 | 1 | 0 | 0 | 0 |
| 25 | 2 | 1 | 1 | 1 | 1 | 0 | 0 |
| 30 | 2 | 1 | 1 | 1 | 1 | 0 | 0 |
| 35 | 0 | 1 | 1 | 1 | 1 | 1 | 0 |
| 40 | 0 | 1 | 1 | 1 | 1 | 1 | 0 |
| 45 | 0 | 1 | 1 | 1 | 1 | 1 | 0 |
| 50 | 0 | 1 | 1 | 1 | 1 | 1 | 0 |
| 55 | 0 | 1 | 1 | 1 | 1 | 1 | 0 |
| 60 | 0 | 1 | 1 | 1 | 1 | 1 | 0 |
| 65 | 0 | 0 | 1 | 1 | 1 | 1 | 2 |
| 70 | 0 | 0 | 1 | 1 | 1 | 1 | 2 |
| 75 | 0 | 0 | 0 | 1 | 2 | 2 | 2 |
| 80 | 0 | 0 | 0 | 1 | 2 | 2 | 3 |
| 85 | 0 | 0 | 0 | 1 | 2 | 2 | 3 |
| 90 | 0 | 0 | 0 | 1 | 2 | 3 | 4 |
| 100 | 0 | 0 | 0 | 1 | 2 | 3 | 4 |

Table S1: Choice of the number of mutator subpopulations with specific mutation rate multiples from $F_{\text{vals}}$ for different biases, $k$, towards larger mutation rate increases.

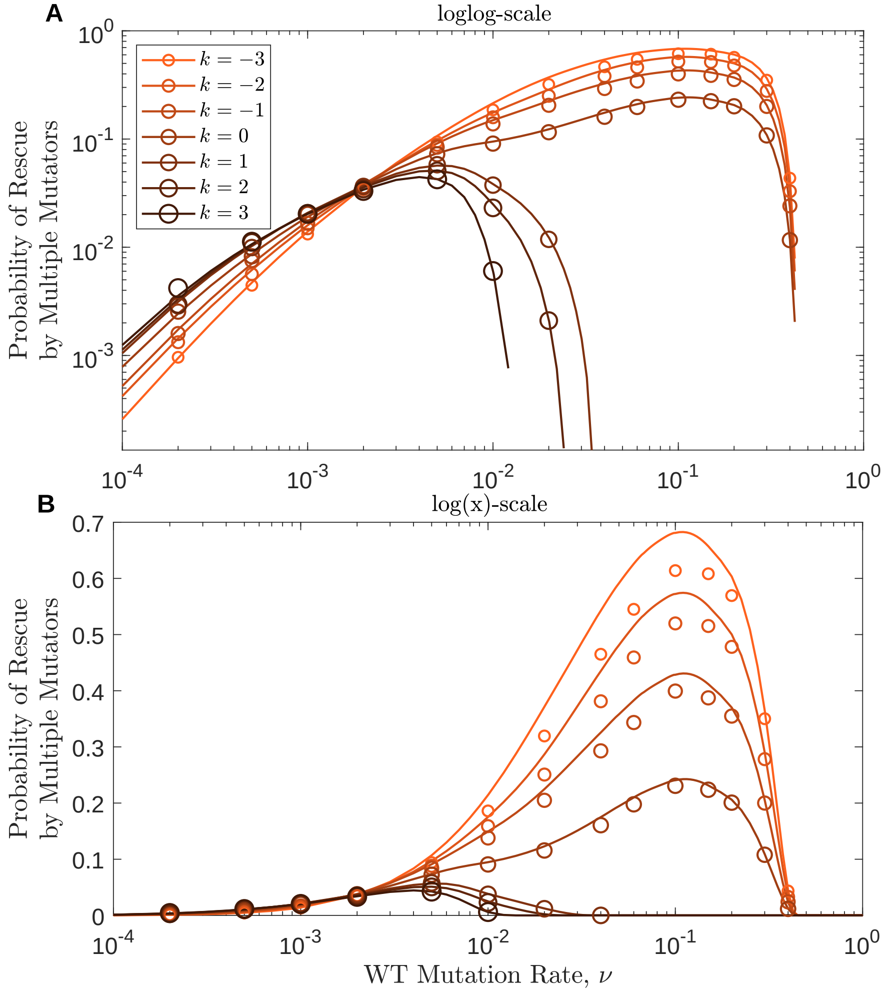


Figure S10: Elevating mutation rates from low wild-type levels incurs minimal costs, thus favouring larger mutation rate multipliers, while this effect flips for high wildtype mutation rates. Twenty mutators emerge with an overall emergence factor $e = 2000$ with different biases towards larger mutation rate increases, $k$. The environmental change speed is $n = 2$ in all cases. Results are shown on a loglog-scale in A and on a log(x)-scale in B. Circles show simulation results of over $5 \times 10^4$.

Gillespie, D. T. 1977. Exact stochastic simulation of coupled chemical reactions. The journal of physical chemistry 81:2340–2361.

Marrec, L., and A.-F. Bitbol. 2020. Adapt or perish: Evolutionary rescue in a gradually deteriorating environment. Genetics 216:573–583.
